# Coupled Mutual Inhibition and Mutual activation motifs as tools for cell-fate control

**DOI:** 10.1101/2022.05.27.493756

**Authors:** Burhanuddin Sabuwala, Kishore Hari, S V Abhishek, Mohit Kumar Jolly

## Abstract

Multistability is central to biological systems as it plays a key role in adaptation, evolvability, and differentiation. Multistability can be achieved by integrating positive feedback loops into the regulation. The simplest of such feedback loops are a) a mutual inhibition (MI) loop, b) a mutual activation (MA) loop, and, c) self-activation. While it is established that all three motifs can give rise to bistability, the characteristic differences in the bistability exhibited by each of these motifs is relatively less understood. Here, we use ODE-based simulations across a large ensemble of parameter sets and initial conditions to study the characteristics of the bistability of these motifs and their limits in the parameter space. We also examine the behavior of these motifs under mutual degradation and self-activation. Finally, we investigate the utility of these motifs for achieving coordinated expression through cyclic and parallel coupling. Through our analysis, we found that MI-based architectures offer robustness in maintaining distinct multistability and allow for coordination among multiple genes. This coordination paves way for understanding the naturally occurring gene regulatory network and may also help in the better design of robust and controllable synthetic networks.

## 1 Introduction

Multistability is an important aspect of multiple biological networks. Two processes that shape the survival and development of organisms - evolvability and differentiation - benefit greatly from multistability^(37)^. Given the stochastic nature of the environments that biological systems (from cells to organisms) are exposed to, multistability can enable efficient adaptation, thus increasing the ‘fitness’ of the system by efficiently responding to changes in the environment^(1,39)^. Multistability also allows for non-genetic heterogeneity in a population, thus improving the chances of the population’s survival during a fitness-based selection in fluctuating environments, as earlier observed in the microbial populations and increasingly reported in multicellular contexts too^(54,22,13,50,38)^. The presence of multistability also allows for multicellular organisms to develop multiple cells/organs with specialized functions, derived from a small set of undifferentiated cells (stem cells)^(25)^. Such functional distribution is a fundamental aspect of mammalian physiology during development, tissue injury as well as cellular reprogramming^(51,2)^. Thus, the investigation of multistability is hence crucial for not only understanding diverse biological systems but also for creating and regulating these different functions via synthetic biology^(60,21,58,57)^.

Over the last few decades, the origins of multistability in biochemical networks have been well-studied through extensive experimental and computational analysis (both deterministic and stochastic simulations)^(44,14,16,46,48,23)^. The necessary condition for multistability is the presence of one or more positive feedback loop(s) that can amplify a system’s property (e.g., gene expression level) after it crosses a certain threshold, thereby creating distinct “high” and “low” states for the said property^(19,3,26)^. The simplest of such feedback loops that give rise to bistability are a) a self-activation (SA) involving a single biological entity (protein/RNA etc) enhancing its own activation (positive auto-regulation)^(17,30,56)^, b) a mutual inhibition loop (MI) where 2 entities inhibit the activation/production of each other, thereby forming an indirect positive feedback loop^(43,52,5)^, and c) a mutual activation loop (MA) where 2 entities activate/stimulate each other^(8)^. While MI motifs have been observed more commonly in decision-making systems such as cellular differentiation, MA motifs are commonly observed in signal amplification^(24)^. The SA motif is often seen in combination with either of these motifs, enhancing the multistability of the system^(18,41,36)^. These motifs are often present in complex gene regulatory networks (GRNs) to control various cell-fate decisions in biological systems.

While all three of these motifs can give rise to multistability, the presence of these motifs themselves is not sufficient for multistability. The appropriate sets of parameters that govern various interactions among these motifs are crucial for the emergence of multistability^(32,7,27)^. Furthermore, the noisy nature of biological systems can alter the likelihood of achieving multistability using these motifs^(33,34,53)^. This becomes especially important in synthetic biology-related studies that aim to introduce these motifs in biological systems to achieve precise control of multistability. A detailed understanding of the behavior of multistability in presence of variability in parameters is lacking in the field. Furthermore, while all three motifs can give rise to bistability, the difference between the dynamical nature of bistability given by each of these motifs is relatively less studied.

Here, we investigate the characteristics of bistability given by the two motifs MA and MI over a large set of randomly chosen kinetic parameter ensemble. We study the nature of the distributions of node activation levels in the motifs and their implications on synthetic control aspects. We then include the components of self-activation and mutual degradation in these motifs and study the changes in multistability over the parameter set ensemble. Finally, we investigate the utility of these motifs in achieving a coordinated expression of multiple nodes in a system through the cyclic and parallel coupling of these motifs to identify the design principles of multistability that can lead to biological phenotypes defined by coordinated expression of multiple nodes.

## 2 Results

### 2.1 Mutual inhibition shows two clearly separable clusters of states, as compared to mutual activation

In a two-node system, bistability can be obtained via two kinds of network connections – mutual activation and mutual inhibition^(21,20,4)^. While mutual inhibition (MI) leads to one of the nodes dominating the other (a high-low or 10 steady state) in every steady state, mutual activation (MA) leads to both nodes together having either high (11) or low (00) expression. This behavior is demonstrated using the nullclines generated via a specific symmetric parameter set for both MI and MA (Fig 1). Using RACIPE^(31)^, a parameter agnostic simulation formalism that samples multiple random parameter sets from a pre-defined parameter space, we probed further into the characteristics of multistability given by these motifs.

**Figure 1:**
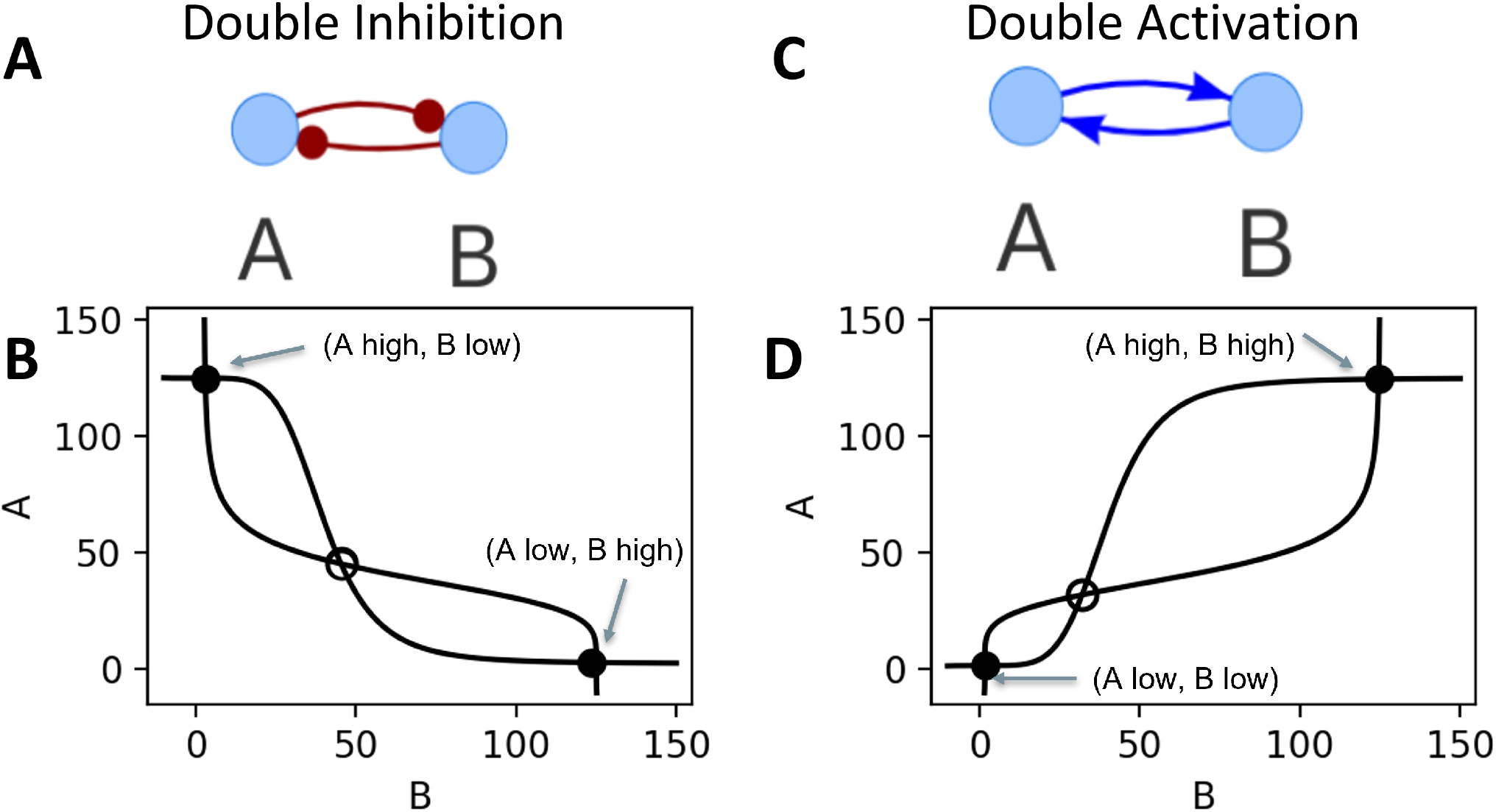
Bistability of Mutual Activation and Mutual Inhibition. a) Network topology of Mutual Inhibition. b) Nullclines for a set of symmetric parameters of Mutual Inhibition (The parameters are given in SI Table1). c) Network topology of Mutual Activation. d) Nullclines for a set of symmetric parameters of Mutual Activation (The parameters are given in SI Table1).

First, we calculated the density distributions of the steady states obtained from these motifs using RACIPE. As expected, in both the motifs, we saw two clear clusters of states, with well-defined and non-overlapping boundaries (Fig 2 A-D). Interestingly, while both the clusters of MI had similar densities and spread (Fig 2 B), the cluster corresponding to low-low state was sparse and had a larger spread than that of high-high state in MA (Fig 2 D, notice the reduced color density). To better quantify this claim, we obtained the distribution of the state-space along the principle component axis (Fig S1B, I). We notice a prominent symmetry in the distribution of PC1 for MI as compared to MA. While MI and MA both can give rise to bistability, addition of self-activations to these motifs can give rise to a greater number of steady states^(32,24,11)^. We asked if the addition of self-activation also changes the nature of the clusters seen in RACIPE. As a control case, we simulated the topologies after adding self-inhibition to both the nodes as well. As expected, the fraction of parameter sets having bistable solutions increased in MA and MI with self-activation (MASA, MISA) as compared to MA and MI, while that of MA and MI with self-inhibition (MASI, MISI) is lower. A small fraction of parameters in MASA and MISA also gave rise to tristability (Fig 2 E,F)^(35,8)^. The steady state clusters in bistable solutions were still distinct and had similar shapes as in MI and MA, but the clusters start to merge in both MISA (Fig 2I, S1D) and MASA (Fig 2J, S1K). The tristable solutions showed interesting patterns. In case of all the tristable solutions of MISA (Fig 2K), we see an additional third cluster emerging. In MASA, we observed an increase in the density of the low-low state (Fig 2L). Furthermore, the third state appears between the first two states, with two distinct densities. The clusters in MASA are not as distinct and discrete as in MISA (Fig **??**E vs L). This implies that even for a large set of parameters, MISA offers a much more discrete control over the stable steady states, while MASA offers a much more continuous control.

**Figure 2:**
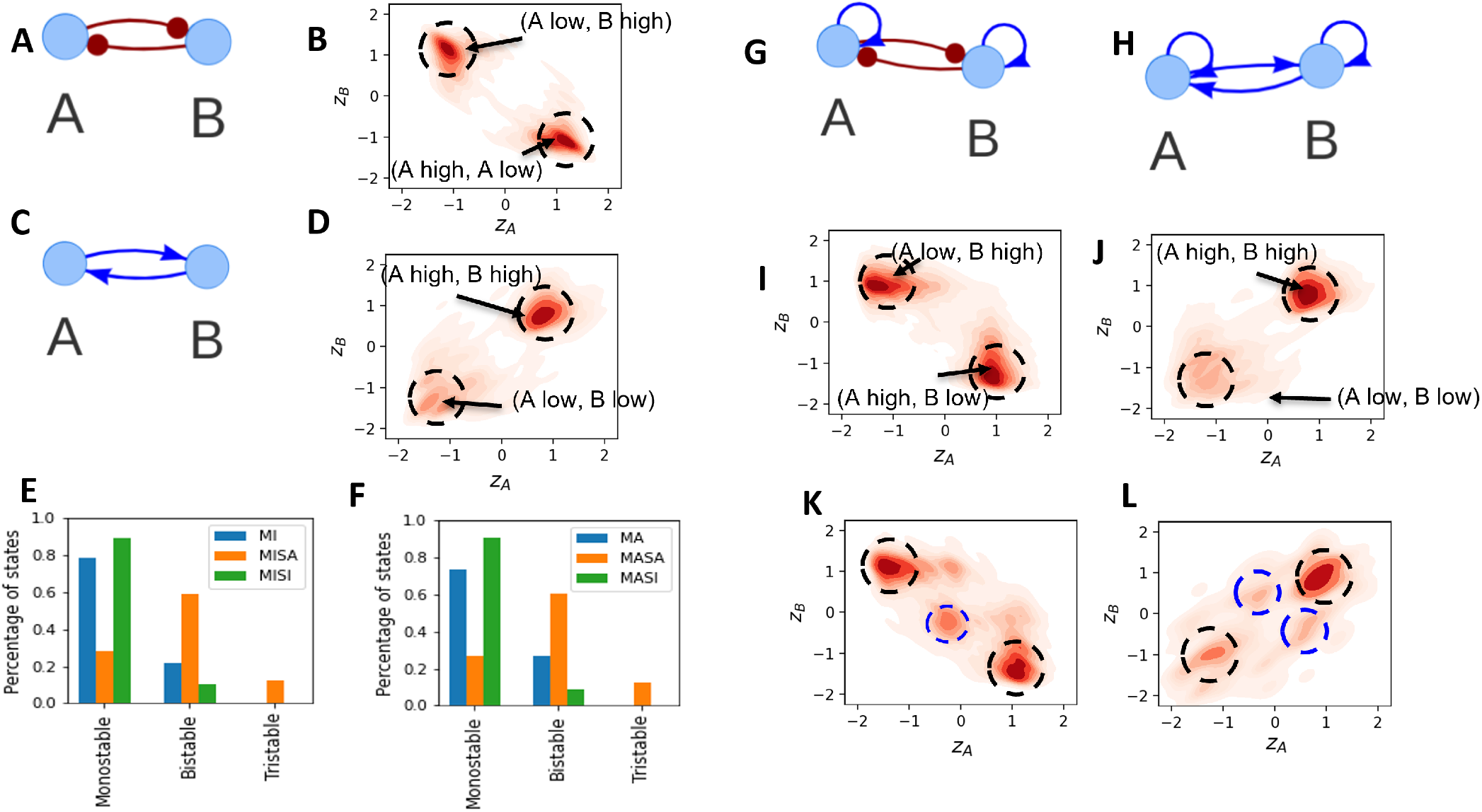
Multistability of Mutual Inhibition (MI) and Mutual Activation (MA) A) Network Topology of MI. B) Density plot depicting all the bistable steady states of MI. Dark color implies higher density of states. Clusters can be seen as boundaries around the dark color densities. C) Network Topology of MA. D) Density plot depicting all the bistable steady states of MA. E) Distribution of a number of steady states for each MI based networks. F) Distribution of the number of steady states for each MA based network. G) Network Topology of MI with self-activation (MISA). H) Network Topology of MA with self-activation (MASA) I) All the bistable steady states of MISA. Two distinct clusters are clearly seen. J) All the bistable steady states of MASA. Two distinct clusters are clearly seen. K) All the tristable steady states of MASA. A third cluster along with the two clusters is seen. L) All the tristable steady states of MASA. No additional cluster is seen apart from the two dominating clusters. There is an enrichment in steady states lying between the two dominating clusters.

We also examined the addition of self-inhibition on both the nodes of mutual activation and mutual inhibition (Fig S2). We found that addition of self-inhibition increase the frequency of parameter sets enabling monostable states at the expense of bistable states, as compared to only mutual activation (MA) or mutual inhibition (MI). Additionally, bistable steady states are also now closer to each other, thereby reducing the separation. Therefore, addition of self-inhibition can reduce the robust controllability expected in a switch-like behavior.

### 2.2 Mutual Degradation has varying effects on dynamics of mutual inhibition and mutual activation motifs

While cooperativity (hill coefficient > 1) is sufficient to achieve multistability in toggle switch^(21)^, multistability can also be achieved in absence of cooperativity via mutual degradation, i.e., the protein products of A and B form an irreversible complex which can no longer activate or inhibit A or B (for instance, it can be marked for degradation) (Fig 3 A,B)^(35)^. Such mechanisms are seen in multiple biological scenarios, including the cell-fate decision circuits in differentiation and development^(49)^. We characterized the dynamics of steady states for varying degrees of mutual degradation for MI and MA.

**Figure 3:**
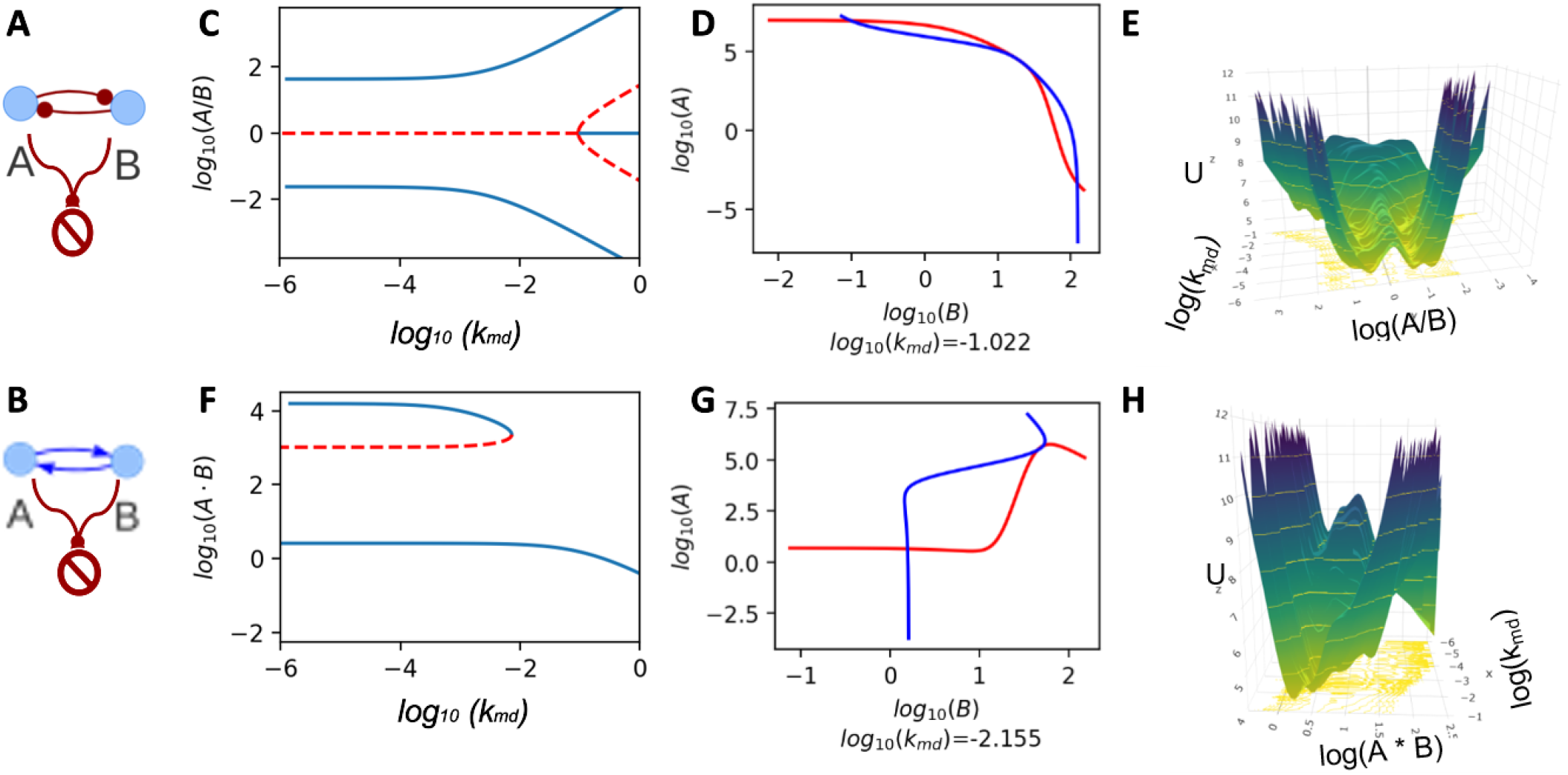
Mutual degradation effects on MI and MA. A) Network topology of MI with mutual degradation. B) Network topology of MA with mutual degradation. C) Bifurcation diagram of MI with mutual degradation with the mutual degradation rate constant *K_md_* as the bifurcation parameter. D) Nullcline plots at the pitchfork bifurcation point. E) Pseudo Potential landscape of MI with mutual degradation. F) Bifurcation diagram of MA with mutual degradation with *K_md_* G) Nullcline plots at the saddle-node bifurcation point. F) Pseudo Potential landscape of MA with mutual degradation.

To understand the effect of mutual degradation on MI and MA network behaviour, we simulated the mutual degradation system at a symmetric set of parameters, i.e., with equal production rate and degradation rate of A and B, and identical parameters for shifted Hill functions depicting the influence of A on B and vice versa (see Methods section, Table 2). We then monitored the steady state dynamics using nullcline and bifurcation analysis. Bistability in MI is not affected for low levels of mutual degradation, as evident by the bifurcation diagram. As the degradation rate increases, the stable steady states start to diverge from one another along the A/B axis (Fig 3C). The nullclines around the bifurcation point (*log*_10_(*k_md_*) ≈ −1.022) demonstrate the emergence of the new state via the tangential interaction of the A and B nullclines at the bifurcation point (Fig 3D, S3B-D). This effect is better visualized in the pseudo-potential landscape of the state-space, generated using a pseudo-potential on the z-axis calculated as negative log probability of occurrence of a given state during stochastic dynamics of the network^(55,40)^ (see Methods, Fig 3E). At larger values of *k_md_*, the unstable state undergoes a pitchfork bifurcation and results in an additional steady state that has the configuration of A low B low, uncharacteristic of an MI circuit (Fig S3A). The emergence of this state is demonstrated in the nullcline at the bifurcation point (Fig 3D; *log*_10_(*k_md_*) ≈ −1.022). In case of MA, as the mutual degradation gets stronger, the high-high stable state collapses along with the corresponding unstable steady state, forming a saddle node (Fig3F,H). The nullclines at the point of bifurcation have been shown to demonstrate how the two states vanish by separation of the two nullclines at the high-high state (Fig 3G, S3F-H, *log*_10_(*k_md_*) ≈ −2.155). Both the nullclines touch each other tangentially at the bifurcation point. These, together with the pseudo potential landscape (Fig 3H), demonstrate the loss of the high-high steady state in MA in presence of mutual degradation. This analysis suggests that mutual inhibition motif is robust in maintaining its switch like behavior and the nature of the steady states, under the influence of mutual degradation.

**Table 1:**
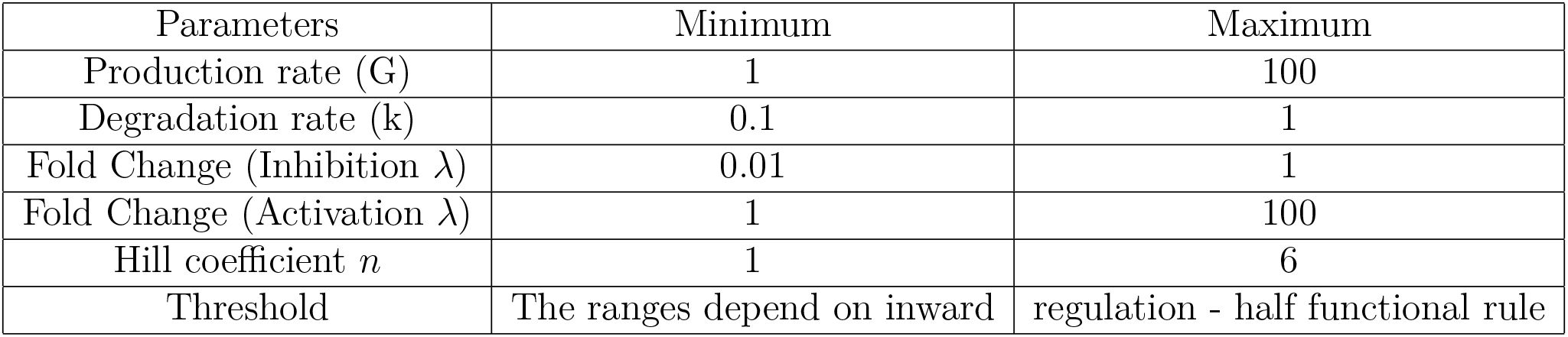
Parameter ranges for RACIPE simulations

**Table 2:** Parameter values for mutual degradation analysis

| Parameters | Mutual Inhibition | Mutual Activation |
| --- | --- | --- |
| Production rate (g) | 50.0 | 50.0 |
| Degradation rate (k) | 0.4 | 0.4 |
| Fold Change ($\lambda$) | 78.0 | 0.02 |
| Hill coefficient ($n$) | 5 | 5 |
| Threshold | 40.0 | 40.0 |

### 2.3 Cyclic Chain architecture for coordinated expression of multiple nodes

While designing electrical circuits, circuit elements are often used in series and parallel configurations to achieve complex behaviors. Similarly in synthetic biology, one might achieve complex and robust biological behaviors if right combinations of switches are used^(9,45)^. One such complex behavior of particular interest in the context of cell-fate determination is coordinated expression of biochemical molecules that stabilize a phenotype. Given that MI and MA circuits give rise to robust phenotypes over a large parameter regime, we wanted to see if architectures constructed using these motifs can achieve coordinated expression of multiple nodes.

The first configuration, the cyclic connection, is formed by connecting multiple mutual activation or mutual inhibition motifs in series in a cyclic fashion, i.e., the last node in the series is connected to the first node. The network obtained by mutual activation (or mutual inhibition) based cyclic connection is named MA (or MI) cyclic chain. Self-activation or self-inhibition could be added on all the nodes to obtain similar variations too. We constructed MI and MA cyclic chains with 10 nodes for our analysis (Fig 4A and 4B). In this configuration, we define the distance between any pair of nodes as the shortest number of edges that the two nodes are separated by, so that neighboring nodes will have a distance of one and so on. Some examples are shown in Fig 4A.

**Figure 4:**
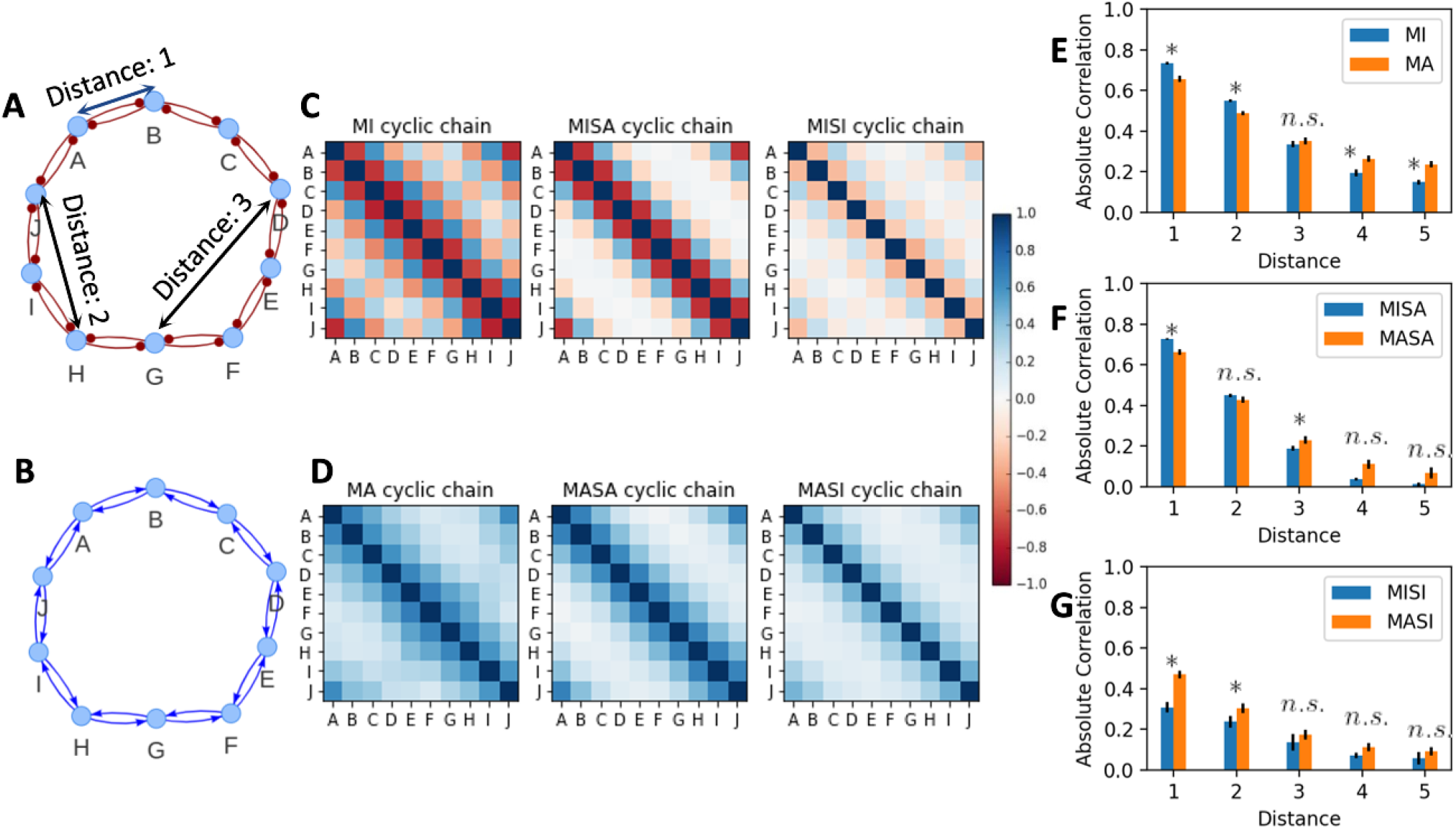
Comparison of Coordinated expression in MI and MA cyclic chains. A) Network topology representation of MI based cyclic chain network. Examples of distance between nodes of the network is shown. B) Network topology representation of MA based cyclic chain network. C) Heat Maps of correlation matrices of all steady states obtained from RACIPE for MI, MISA and MISI cyclic chains from left to right. D) Heat Maps of correlation matrices of all steady states obtained from RACIPE for MA, MASA and MASI cyclic chains from left to right. The colorbar key for both C and D panels is shown. E) Variation of the absolute value of correlation between the nodes at a given distance for MI and MA networks is shown. F) Variation of the absolute value of correlation between the nodes at a given distance for MISA and MASA networks is shown. G) Variation of the absolute value of correlation between the nodes at a given distance for MISI and MASI networks is shown. [* – *p* < 0.001 and * * – *p* < 0.0001]

Pairwise correlation heatmaps of the normalized RACIPE solutions for these cyclic networks reveal that mutual inhibition-based networks show a chessboard-like correlation behavior, i.e., pairs of nodes separated by odd distance (distance = 1 or 3 or 5, ex: A-B, A-D, B-C etc) show a negative correlation (red color), while pairs separated by even distance (distance = 0 or2 or 4, ex: A-C, B-D etc) show a positive correlation (blue color)(Fig 4C). Addition of self-activation to the nodes of MI cyclic chain does not disrupt this pattern. The strength of the correlation in MISA cyclic chain, however shows a sharp decrease after a distance of 2, as compared to that of MI cyclic chain (Fig S4A, notice the white patches in the MISA cyclic chain heatmap in Fig 4C). MISI cyclic chain on the other hand shows much weaker correlation strength than MI and MISA cyclic chains. Comparing MI, MISA and MISI cyclic chains, one can infer that MI cyclic chain is a better candidate for coordinated expression (Fig S4A). In MA cyclic chains, the correlation between any pair of nodes remains positive (Fig 4D). The loss of correlation strength upon addition of SA and SI to the nodes of MA cyclic chain is observed here as well (Fig S4B, Fig S5).

We then compared the strengths of correlations obtained by MI vs MA cyclic chains. We find that, while MI cyclic chain shows stronger correlation for nodes separated by shorted distance, the strength of correlation decays faster with node distance than that of MA cyclic chain, leading to the later showing stronger correlations at longer distance between nodes (Fig 4E). Similar trend is observed upon the addition of self activation, however the difference at longer distances becomes insignificant (Fig 4F). Upon adding self-inhibition, however, MASI cyclic chain shows stronger correlations at all distances in comparison to MISI cyclic chain (Fig 4G).

Together, these results suggest that at shorter distances, MI cyclic chain performs better than MA cyclic chain. While MA performs better for longer distances, the strength of correlation at longer distances is low. Hence, we moved on to the parallel architecture, where the distance between the nodes remains short, as will be explained in the following section.

### 2.4 Parallel architecture provides better coordination than cyclic architecture

The parallel configuration consists of a central node connected to the peripheral nodes by either MI (MI parallel core) or MA (MA parallel core) motif, such that the central node is in parallel influencing all the peripheral nodes. For our analysis, we generated a network of 10 peripheral nodes which are connected with a central node, making the total nodes in the network 11 (fig 5 A,B). Additionally we also considered the networks in which self activation (MISA parallel core, MASA parallel core) or self inhibition (MISI parallel core, MISA parallel core) is added on all the 11 nodes.

**Figure 5:**
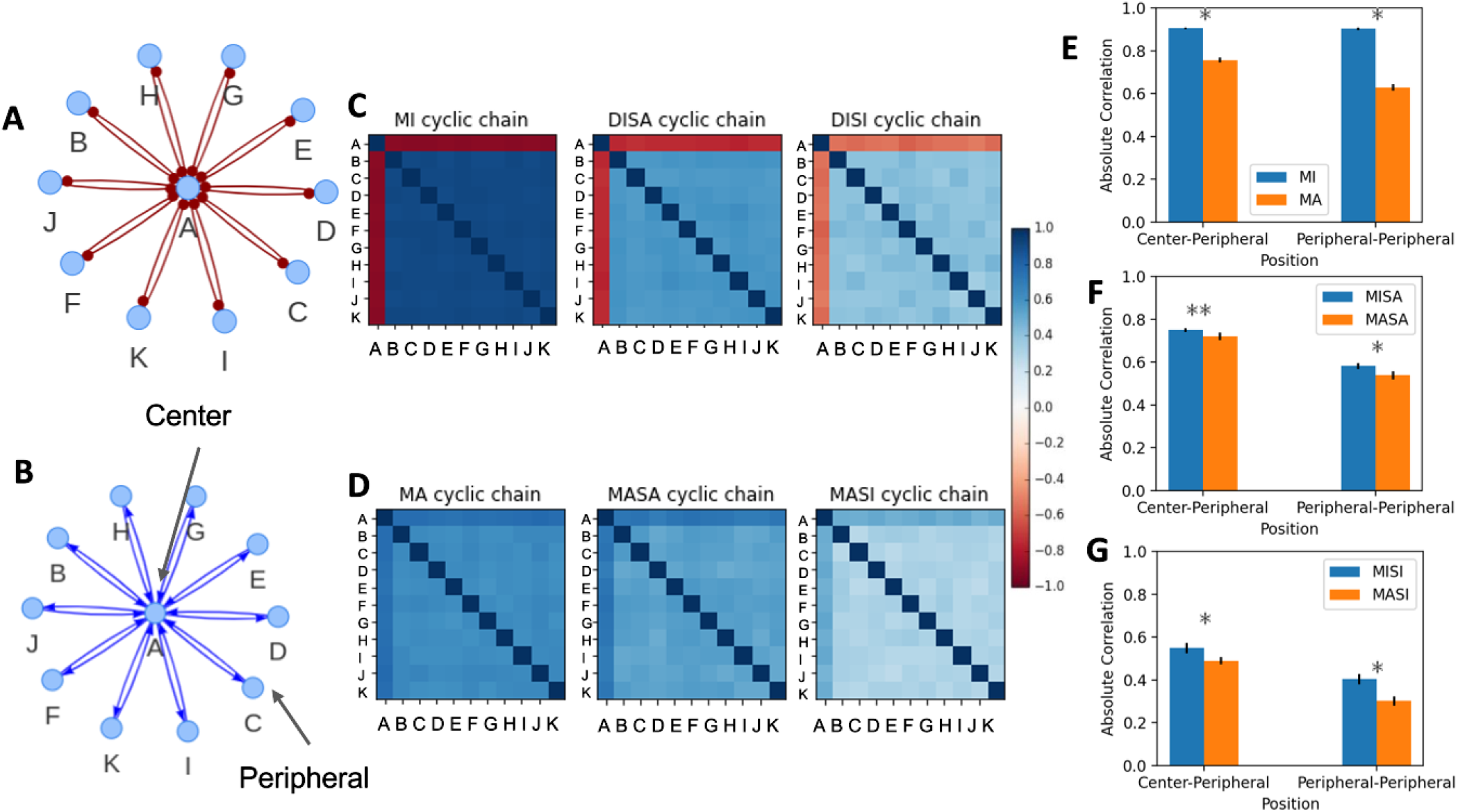
Comparison of Coordinated expression in parallel cores constructed using MI and MA motifs. A) Network topology representation of MI based parallel network. B) Network topology representation of MA based parallel network. Center and peripheral nodes are labelled. C) Heat Maps of correlation matrices of all steady states obtained from RACIPE for MI, MISA and MISI parallel cores from left to right. D) Heat Maps of correlation matrices of all steady states obtained from RACIPE for MA, MASA and MASI parallel cores from left to right. E) Comparison of the absolute correlation between central-peripheral and peripheral-peripheral for MI and MA parallel cores. F) Comparison of the absolute correlation between central-peripheral and peripheral-peripheral for MISA and MASA parallel cores.g) Comparison of the absolute correlation between central-peripheral and peripheral-peripheral for MISI and MASI parallel cores.

In MI parallel core, the central node shows a negative correlation with all peripheral nodes, while the peripheral nodes all show a positive correlation with each other (Fig 5C). Addition of SA and SI to the nodes of parallel core resulted in a reduction of both central-peripheral (C-P) as well as peripheral-peripheral (P-P) node correlation (Fig S6A). In MA parallel core, all correlations are positive, with center-peripheral correlation being stronger than P-P correlation (Fig 5D). Similar to the MI based core, addition of SA and SI to the nodes of MA parallel core showed a decreased correlation strength (Fig S6B).

The P-P node correlations in parallel cores are a measure of the coordinated expression of the peripheral nodes. To understand which motif (MI vs MA) performs better in achieving coordinated expression, we compared the mean strength of correlation between C-P nodes as well as P-P nodes in MI parallel core and MA parallel core circuits (Fig 5E). C-P correlation is the same as correlation between nodes forming MI/MA motifs. Clearly, the strength of correlation is higher for MI than for MA. MI parallel core’s P-P correlation also showed a higher mean than that of MA parallel core. Furthermore, the C-P and P-P correlations for MI parallel core showed similar mean strengths, while P-P correlations for MA parallel core were weaker than C-P correlations. This behaviour can be explained by the high level of heterogeneity observed in MA steady states (Fig 2B,D). Furthermore, MI parallel core had significatly higher C-P and P-P correlation strengths than MA parallel core despite the addition of SA and SI to the nodes (fig 5F,G), thus showcasing better coordination in MI circuits relative to MA ones.

### 2.5 Combining Parallel core and cyclic chains for enhanced coordination

Both parallel and cyclic conformations enable coordinated expression of the nodes in the network. We then asked if combining parallel and cyclic conformations increases the synchronization. To answer this question, we generated a set of five networks and compared their dynamics: i) A parallel core (core), ii) addition of a single ring-like connection covering all the nodes (core + 1 ring), iii) addition of mutual regulatory ring-like connection covering all the nodes (core + 2 rings / cycle + mutual), iv) a cyclic chain with a central node connected to all the nodes via single activation or inhibition links (cycle + C-P) and v) a cyclic chain (cycle). We considered the networks consisting of 10 peripheral nodes, which are part of the ring and connected to their two immediate neighbors and 1 central node, the which is not the part of the ring and it is connected to all the peripheral nodes. The rings and cores could be either inhibitory or activatory in nature. While MI cycle shows stronger P-P correlation than MA cycle, addition of MI cycle to MI/MA core would cause frustration (since neighbour peripheral nodes are connected by MI motif making them antagonistic, but also controlled by the central node in a similar fashion making them co-express). Hence, we chose to add MA cycle to MA and MI cores to study the coordinated expression of peripheral nodes (Fig 6A, **??**A). To study the extent of synchronization of the peripheral nodes, we employed the calculation of the absolute correlation for different pairs of nodes. Since the correlation in cyclic architecture depends upon the distance between the nodes, we defined PP-n correlations, where PP stands for peripheral-peripheral and n is the distance between the given pair of peripheral nodes.

**Figure 6:**
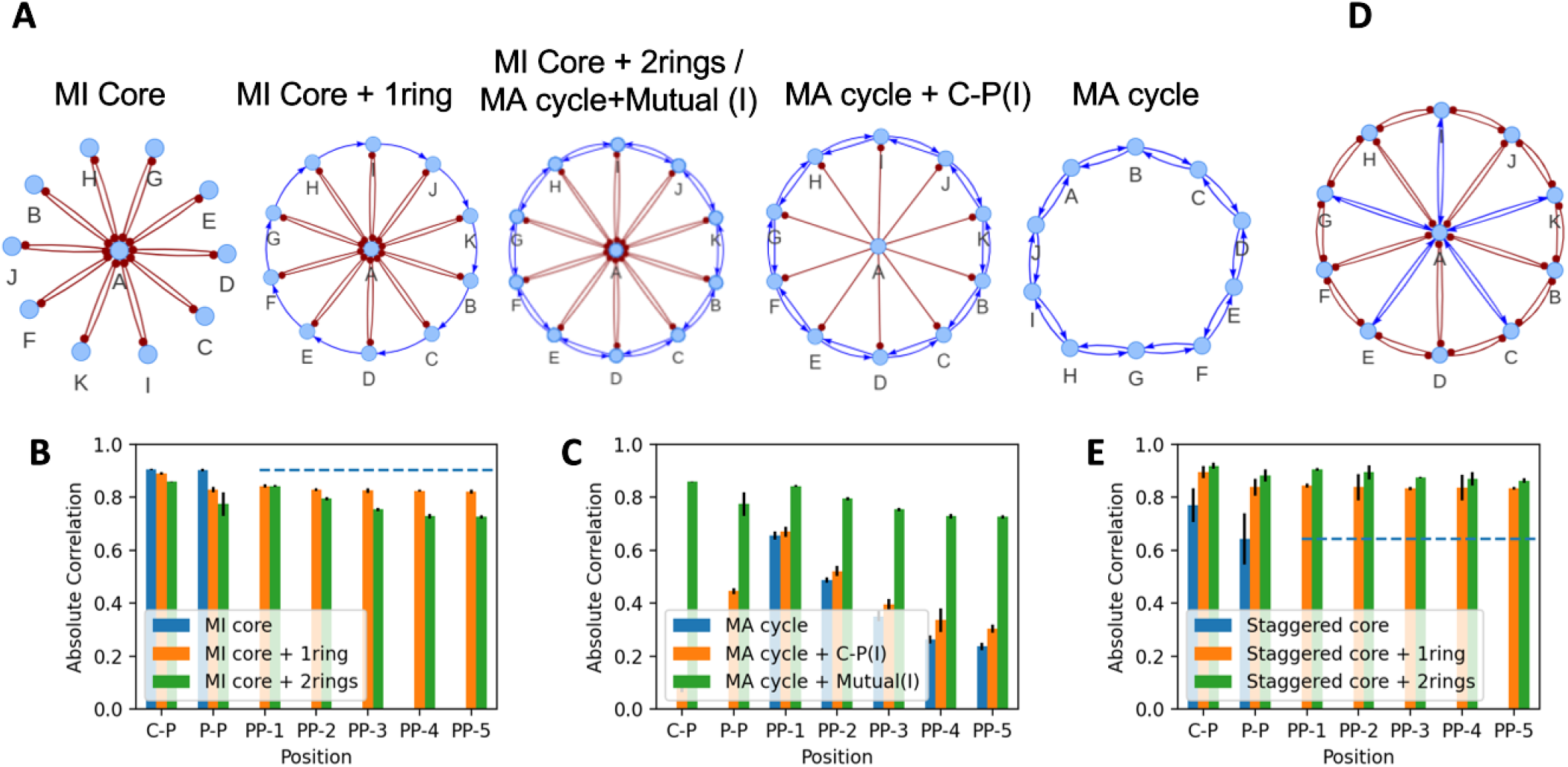
Combinations of cyclic and core architectures for enhanced coordination. A) Five networks designed to systematically test the effect of the combination of an inhibiting core and an activating ring on coordinated expression of the peripheral nodes. B) Comparison of absolute correlation between central-peripheral and peripheral-peripheral nodes at various distances for MI core without a ring, with 1 ring and 2 rings (cycle). Dotted blue line indicates the PP-n correlation strength for MI core. C) Comparison of absolute correlation between central-peripheral and peripheralperipheral nodes at various distances with different degrees of connection with the central node. D) Staggered Core with MI ring network topology. E) Comparison of absolute correlation between central-peripheral and peripheralperipheral nodes at various distances in the staggered core with different degrees of connection among the peripheral nodes. Dotted blue line indicates the PP-n correlation strength for MI core.

We first looked at addition of MA cycle to MI core (Fig 6B). The C-P correlation for MI core was slightly higher than MI core + 1 ring and MI core + 2 rings, suggesting that addition of rings to MI core weakens the effect of MI core on the peripheral nodes. The difference was amplified in P-P correlation, where the correlation strength was significantly higher for MI core in comparison to the addition of 1 and two rings. We then looked at PP-n correlations. In cyclic chains, the correlation between nodes decreased as the distance increased. While a decrease is still seen in MI core + 2 rings, the decrease is much smaller as compared to MA cycle alone (0.75 in MI core + 2 rings vs 0.25 for MA cycle in Fig 4E). MI core + 1 ring showed better resilience to distance between the nodes, but was lesser than the P-P correlation for MI core (dotted blue line in Fig 6B). As expected, MA cycle performs poorly in comparison to MI core + MA cycle (Fig 6C). Addition on single inhibition from central to peripheral nodes shows poor improvement in absolute correlation. Addition of MA cycle to MA core on the other hand shows improved absolute correlation between peripheral nodes as compared to MA cycle alone and MA core alone (Fig S7B). Furthermore, addition of MA core to MA cycle increases the PP-n correlations and makes them robust to the increase in distance. However, the maximum P-P correlation in MA core + 2 rings is lesser than that obtained from MI core (0.85 vs 0.9), making MI core the better architecture for achieving coordinated expression o peripheral nodes.

We then wanted to check if a non-frustrating addition of a core to MI cycle can achieve better coordination in peripheral nodes, since MI cycle achieved better coordination and MA cycle and addition of core to MA cycle showed enhanced coordination between peripheral nodes in all scenarios. Hence, we constructed a staggered core, where the central node connects with peripheral nodes alternatively with MA and MI motifs (Fig 6D). The staggered core without the MI ring showed weak correlations both in C-P and P-P (Fig 6E, blue bars and blue dotted line). However, addition of the inhibitory ring considerably improved the C-P, P-P and PP-n correlations in the network. Addition of 2 rings to staggered core (MI cycle + staggered core) showed the highest absolute correlations for C-P, P-P and PP-n (mean > 0.9 in all cases). Hence, we find that MI cycle + staggered core offers the best possible coordinated peripheral node expression, followed by MI core and MA cycle + MA core.

## 3 Discussion

Biological systems are often viewed as complex control systems. By means of regulatory networks, multiple control functions, such as homeostatic systems, oscillatory systems, switching systems etc. are maintained^(21,15,59)^. Networks underlying switching systems are particularly well studied since they play a crucial role in the decision-making involved throughout the course of the development of an organism. While constructing such networks from experimental data and connecting their emergent properties to the observed biological phenotype is the norm, the reverse problem has gained much attention. That is, given a phenotype, puzzling out the network topology and the corresponding parametric regime in which said phenotypes can be observed. There are two reasons for this. First, solving such a problem allows biologists to look for specific interaction patterns in the underlying biology, either via experiments like flow cytometry or in high throughput data like RNA-seq, ChIP-seq, etc. Second, such information is necessary to construct biomolecular circuits that, when inserted into biological systems such as cells, provide the desired function within the desired margin of error.

To pave the way from phenotype to network motifs, we took the approach of studying common network topologies in detail, understanding the differences in the emergent properties of such networks, and predicting the emergent properties of the networks obtained by coupling these motifs. In this study, we show such analysis of two fundamental motifs of multistability in gene regulatory networks (GRNs) – mutual activation (MA) and mutual inhibition (MI). To study these motifs, we used a parameter agnostic approach called RACIPE^(31,32)^. For any given GRN, RACIPE generates an ensemble of possible stable steady states (phenotypes) in a given parameter space, there by mapping the behavior of the network in a large parameter regime. This becomes important from a synthetic biology perspective, since regulating the interaction strengths of a synthetic circuit is not always as feasible as constructing the circuit itself^(6,47)^.

The ensemble obtained from RACIPE is a continuum of states divided into clusters reflecting the qualitative nature of the motifs. The first clear difference between MA and MI motifs was that the MI clusters were much more symmetric than MA clusters. Additionally, the clusters corresponding to each state were well separated as well. In comparison, the MISA and MASA networks had a higher overlap between the clusters. Particularly in the tristable parameter sets for both MISA and MASA, the third steady-state cluster falls between the pre-existing steady states. This indicates that self-activation makes the control by the bistable switch more continuous, in addition to increasing the multistability. Furthermore, MA offers a more continuous control as compared to MI. Such continuous control can be useful in providing the system with higher adaptability but would also make the system susceptible to environmental fluctuations. Therefore, robust systems would likely have and should be designed using MI circuits, while adaptable systems would benefit from MASA.

We further studied the effect of mutual degradation of the nodes involved in the bistable switches, on the emergent bistability. Interestingly, while MI retained the two prominent states (low-high and high-low) and gave rise to a new state under a high mutual degradation rate constant (*K_md_*), MA lost the high-high state and became monostable under high *K_md_*. This result makes intuitive sense. A high concentration of both the nodes leads to a high degradation rate, thereby supporting states that have a low concentration of at least one of the nodes. While both states of MI satisfy this condition, only the low-low state of MA satisfies this condition. From a system control perspective, this indicates that MI provides a robust control in comparison to MA in systems that incorporate mutual degradation. Given the ability of mutual degradation to give rise to bistability in low cooperativity, a toggle switch (MI) with mutual degradation might be an easier way to achieve multistability. Recent studies from Elowitz group employ mutual degradation-based circuits and demonstrated their utility as a multi-fate architecture and scalability^(60)^.

At an individual level, MI seems to offer more robust control over MA. One such architecture recently explored was a serial and parallel connection of positive feedback loops, where the authors showed that a parallel architecture might help in limiting cell-cell variability, and hence is a preferred architecture in differentiation networks^(10)^. In previous work, we had explored a variety of the serial connection, a cyclic chain of MI motifs labelled circular toggle polygons^(29)^. Given the additional feedback embedded in the circular/cyclic connection, it should be a better candidate for a synchronous expression of nodes potentially leading to the stabilization of cell-fates. Similar to the small-scale architectures, parallel and cyclic connections constructed using MI circuits provide higher coordination than those constructed using MA circuits. Between the two architectures, we find that parallel architecture is better for the coordinated expression of multiple nodes as compared to the cyclic architecture. Similar to the small-scale architectures, parallel and cyclic connections constructed using MI circuits provide higher coordination than those constructed using MA circuits.

In a cyclic architecture, the coordination lasts only upto a small number of serially connected nodes. Such behavior is intuitive since as the distance increases, the path length for the propagation of control increases. While this is not desirable for global control, this feature of limited distance control can potentially impart a certain type of modularity to the system. This would differ from traditional modularity in that the modules would not be independent entities but would have overlapping elements^(42)^. Given the recent understanding of how biological phenotypes are stabilized^(28)^, a parallel MI architecture might be preferred in biological networks as well to coordinate multiple molecules together. This could also explain the power-law nature of large-scale biological networks.

One might expect a combination of parallel and cyclic architectures to provide maximum coordination between multiple nodes of the network. Surprisingly, we find that the addition of cyclic architecture to the parallel MI architecture only lessens the correlation between the peripheral nodes. Interestingly, the addition of a staggered parallel architecture to a cyclic MI architecture (to avoid conflicting interactions to the network) improve the coordination of the individual architectures dramatically. This improved coordination equals the coordination provided by parallel MI architecture and does not overcome it. These results reinforce the conclusion that MI provides better control over the expression levels as compared to MA. Furthermore, MI architectures may also be more susceptible to synergistic action with other architectures.

Given the robust control that MI-based architectures exhibit as compared to MA-based architectures, the reason behind the choice of MA circuits for multistability in biological systems is unclear. One possible role could be in systems that need to be sensitive to even slight changes in the environment around them. Our analysis does make it clear, however, that to construct a multistable system, MI-based architectures provide more robustness both in terms of maintaining distinct multistability and in allowing for coordination among many nodes, which is often a requirement of stabilizing phenotypes. These results are also valid for a large-scale parameter regime given the nature of the methods applied.

## 4 Methods

### 4.1 RAndom CIrcuit PErturbation (RACIPE)

RACIPE is a tool for characterizing the dynamics of Gene Regulatory Networks (GRNs)^(31,32)^. It first generates a system of coupled Ordinary Differential Equations (ODE) corresponding to the given GRN. For a node X in the GRN with a set of input nodes *A_i_* and *I_j_* that activate and inhibit X, respectively, the corresponding ODE is given by the equation:

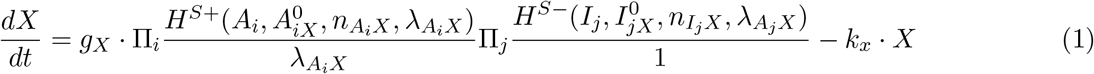

where, *X*, *A_i_*, and *I_j_* are the concentration of the corresponding species. *g_x_* and *k_x_* denote the production rate and degradation rate of species X. 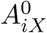 is the half-maximal threshold value of the concentration of *A_i_*, as defined for Hill functions. n is the hill coefficient which characterizes the extent of non-linearity in the regulation of X. λ represents the fold change in the target node (*X*) concentration upon over-expression of regulating node. The functions *H*^*S*+^ and *H*^*S*−^ are known as shifted hill functions^(32)^ and represent the regulation of the target node by the regulatory node.

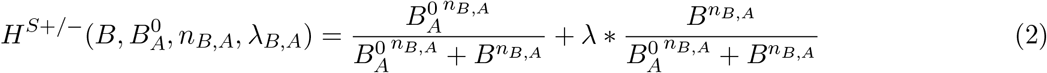

Note that, as the regulatory nodes are overexpressed, *H*^*S*+/-^ approaches. λ > 1 denotes activation; *λ* < 1 denotes repression.

For the system of ODEs generated this way, RACIPE samples parameter sets from pre-defined parameter ranges given below using a uniform distribution.

For each parameter set, RACIPE simulates the system from multiple random initial conditions and obtains the corresponding steady states. For our analysis, we used a sample size of 1000 parameter sets for small networks (MI, MA) and 10000 parameter sets for larger networks (cyclic, parallel and mixed architectures), along with 100 initial conditions for each parameter set. Euler’s method of numerical integration was used for ODE-based simulation. We use the ensemble of steady states obtained for all parameter sets for further analysis.

### 4.2 Normalization and z-score calculation of RACIPE output

To enable better comparison of the steady states across parameters, we normalize the steady state levels as follows:

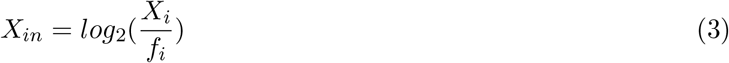

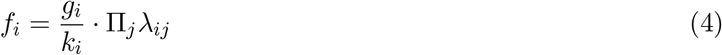

where for the *i^th^* node, *X_i_* is the expression value, *f_i_* is the normalization factor, *g_i_* is the production rate of *i*, *k_j_* is the degradation rate of *i*, and λ*_ij_* is the fold change corresponding to the link from the *j^th^* node to the *i^th^* node.

The normalized expression levels are further converted to z-scores by mean-zeroing and scaling using the standard deviation of the combined normalized expression values.

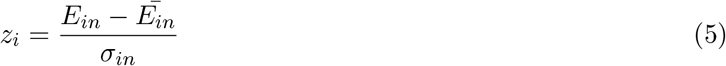

where 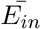 is the combined mean and *σ_in_* is the combined standard deviation.

### 4.3 Mutual Degradation

Mutual degradation is included in the ODE system (equation(1)) by adding the term - *k_md_* · *A* · *B* to each ODE, where *A* and *B* are the concentration of species involved in mutual degradation. The corresponding equations are as follows:

**Mutual Activation with Mutual degradation:**

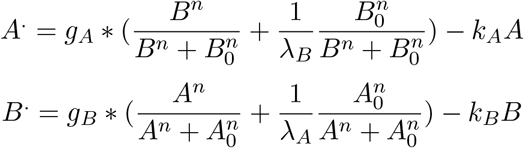
**Mutual Inhibition with Mutual degradation:**

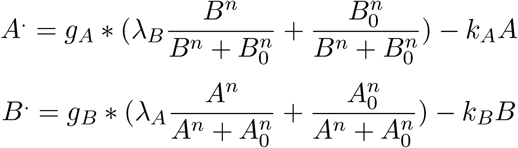

The parameters used to simulate the system are given below:

### 4.4 Nullclines and Bifurcation Plots

For a given ODE system, the nullclines for species *X* are obtained by setting the value of 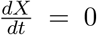 in equation (1), in other words, fixing the concentration of *X* at a certain value. The equation is solved for the corresponding species and the values are plotted. We used the `matcont` package in MATLAB R2018b to obtain the bifurcation plots^(12)^. The parameters used for obtaining the bifurcation plots are given in Table S1.

### 4.5 Calculation of Distance-wise Absolute Correlation

The distance between two nodes is defined by the length of the smallest path in the network between those two nodes. Pearson correlation (calculated using the `corrcoeff` method from `numpy` package in Python 3.7.9) is calculated among every pair of nodes separated by the same distance. The mean and standard deviation of these correlation values is used to compare multiple network configurations.

## 5 Author Contributions

MKJ designed and supervised research; KH, BS and ASV conducted research and analyzed the data; all authors discussed results and participated in the preparation of the manuscript.

## 7 Data and Code Availability

The data generated and analysed for the manuscript can be found at: https://github.com/BurhanSabuwala/DI-DA-robustness

## 8 Acknowledgements

This work was supported by Ramanujan Fellowship awarded to MKJ by Science and Engineering Research Board (SERB), Department of Science and Technology (DST), Government of India (SB/S2/RJN-049/2018), by Infosys Young Investigator award to MKJ supported by Infosys Foundation, Bangalore and by Prime Minister’s Research Fellowship awarded to KH.

## 1 Supplementary Figures

**Figure S1:**
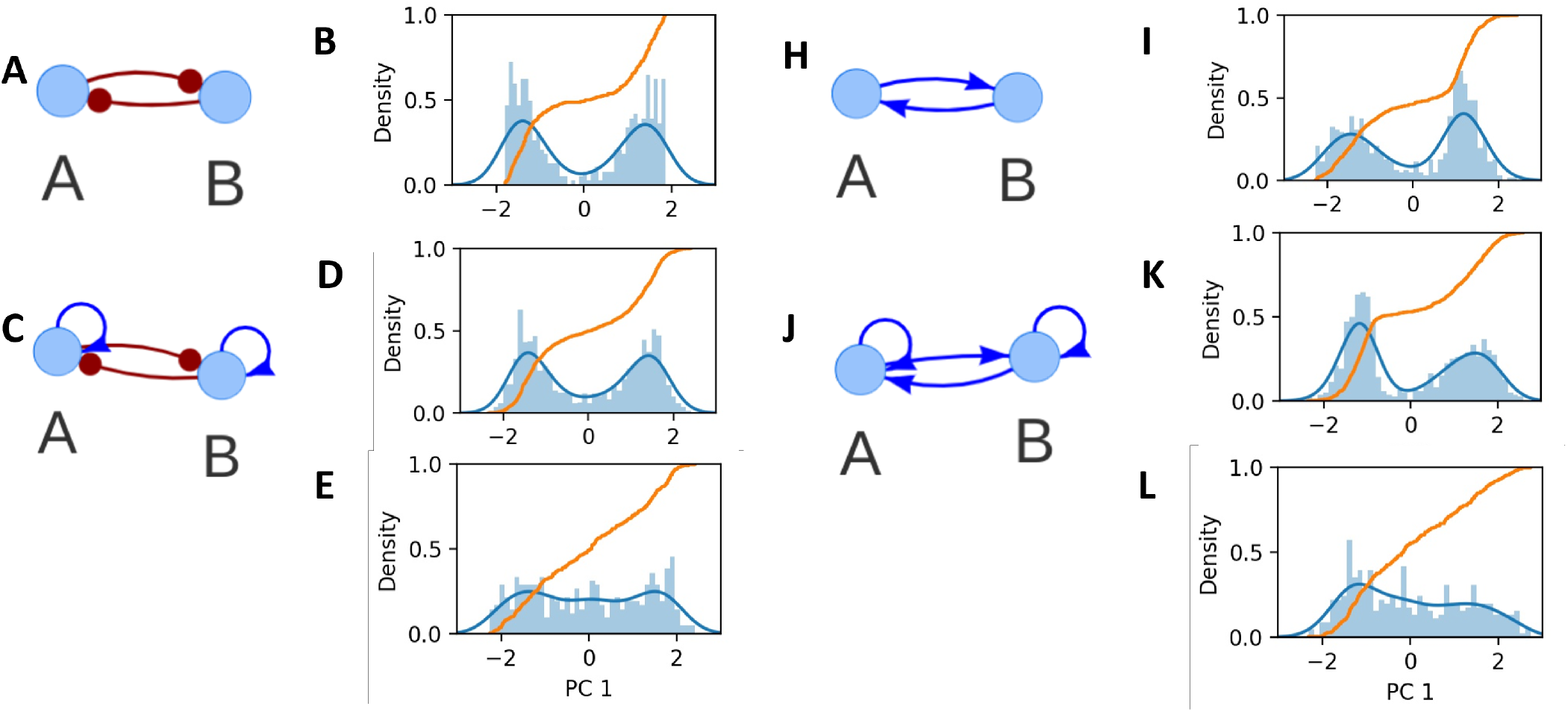
Distribution along the first principle axis of the RACIPE steady states corresponding to the networks. a) Network topology of Mutual Inhibition. b) Distribution of states along the first principle axis of all the bistable steady state solutions of Mutual Inhibition. c) Network topology of Mutual Inhibition with Self Activation. d) Distribution of states along the first principle axis of all the bistable steady state solutions of Mutual Inhibition with self-activation. e) Distribution of states along the first principle axis of all the tristable steady state solutions of Mutual Inhibition with self-activation. f) Network topology of Mutual Inhibition with Self Inhibition. g) Distribution of states along the first principle axis of all the bistable steady state solutions of Mutual Inhibition with self-inhibition. h) Network topology of Mutual Activation. i) Distribution of states along the first principle axis of all the bistable steady state solutions of Mutual Activation. j) Network topology of Mutual Activation with Self Activation. k) Distribution of states along the first principle axis of all the tristable steady state solutions of Mutual Activation with self-activation. l) Distribution of states along the first principle axis of all the tristable steady state solutions of Mutual Activationwith self-activation. m) Network topology of Mutual Activation with Self Inhibition. n) Distribution of states along the first principle axis of all the bistable steady state solutions of Mutual Activation with self-inhibition. The blue line in the plots signify the probability distribution as approximated using gaussian kernel density estimate. The yellow line in the plots is the corresponding cumulative density function.

**Figure S2:**
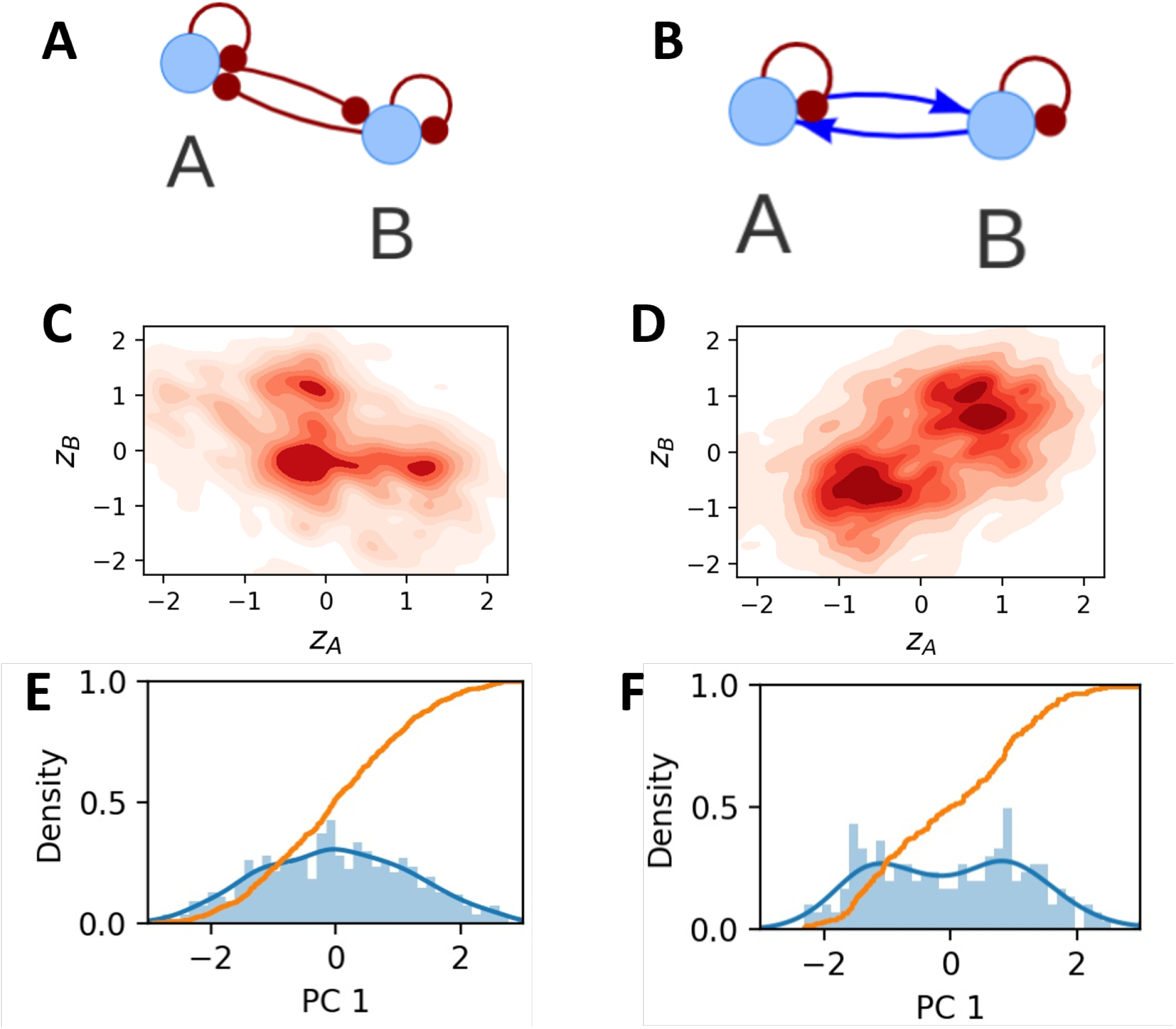
Distribution of states on addition of self-inhibition. a) Network topology of Mutual Inhibition with self inhibition b) All the bistable steady states of Mutual Inhibition with self inhibition. Multiple non-distinct clusters are seen. c) Network topology of Mutual Activation with self inhibition d) All the bistable steady states of Mutual Activation with self inhibition. Multiple non-distinct clusters are seen.

**Figure S3:**
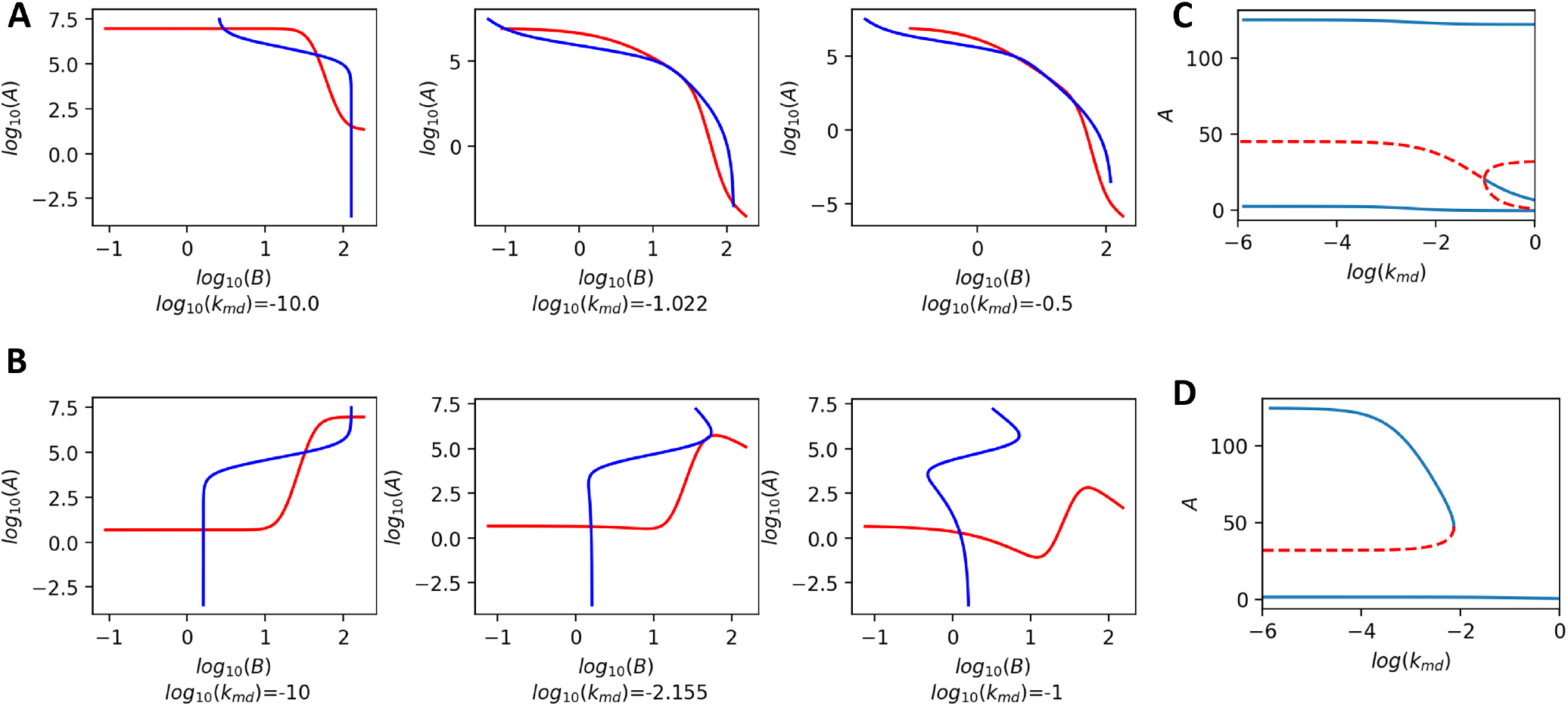
Mutual Degradation. a) Bifurcation diagram of Self Inhibition with Mutual Degradation. b) Nullclines of Self Inhibition with Mutual Degradation at *log*_10_(*k_m_d*) = −10. c) Nullclines of Self Inhibition with Mutual Degradation at log_10_(*k_md_*) = −1.022. This corresponds to the subcritical pitchfork bifurcation point given in fig a). c) Nullclines of Self Inhibition with Mutual Degradation at *log*_10_(*k_md_*) = −0.5. d) Bifurcation diagram of Self Activation with Mutual Degradation. f) Nullclines of Self Activation with Mutual Degradation at *log*_10_(*k_m_d*) = −10. g) Nullclines of Self Activation with Mutual Degradation at *log*_10_(*k_md_*) = −2.155. This corresponds to the saddlenode bifurcation point given in fig e). h) Nullclines of Self Inhibition with mutual degradation at *log*_10_(*k_md_*) = −1

**Figure S4:**
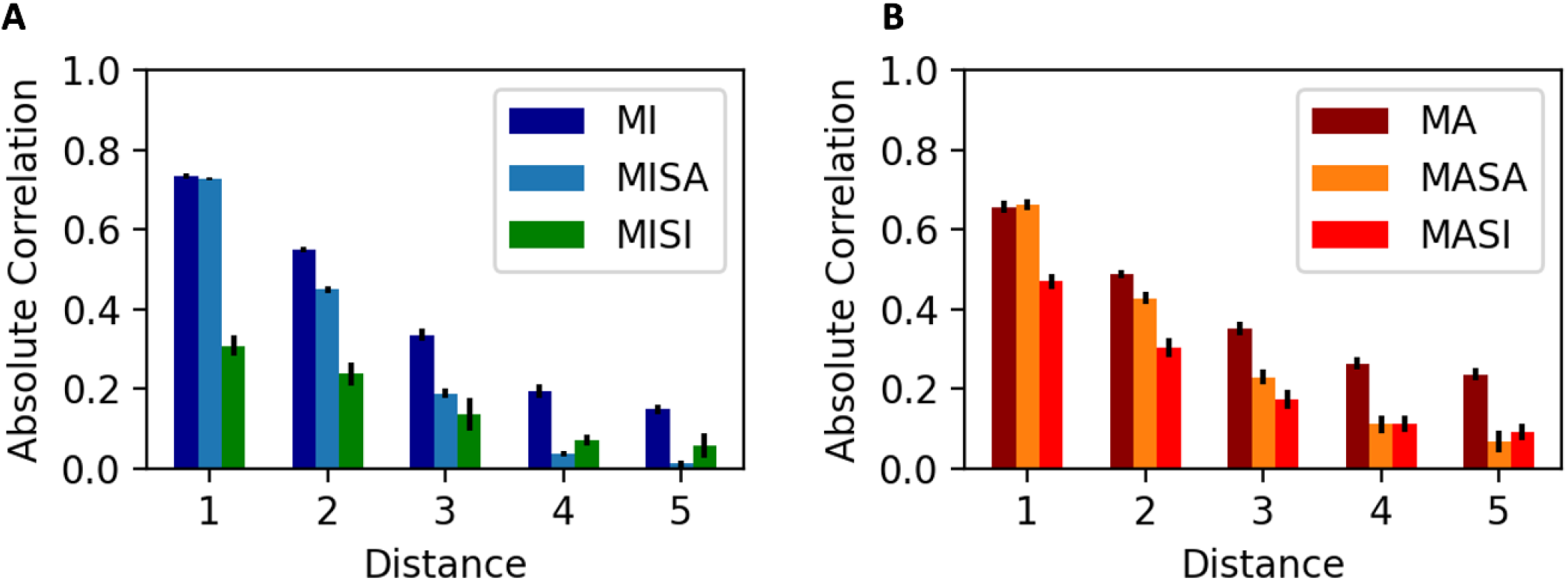
Cyclic Chain Connection. a) Variation of Absolute Correlation with addition of self activation or self inhibition to all the nodes of Mutual Inhibition based cyclic chain connection. The distance along the x-axis is the shortest distance between any two nodes along the chain. b) Variation of Absolute Correlation with addition of self activation or self inhibition to all the nodes of Mutual Activation based cyclic chain connection. The distance along the x-axis is the shortest distance between any two nodes along the chain.

**Figure S5:**
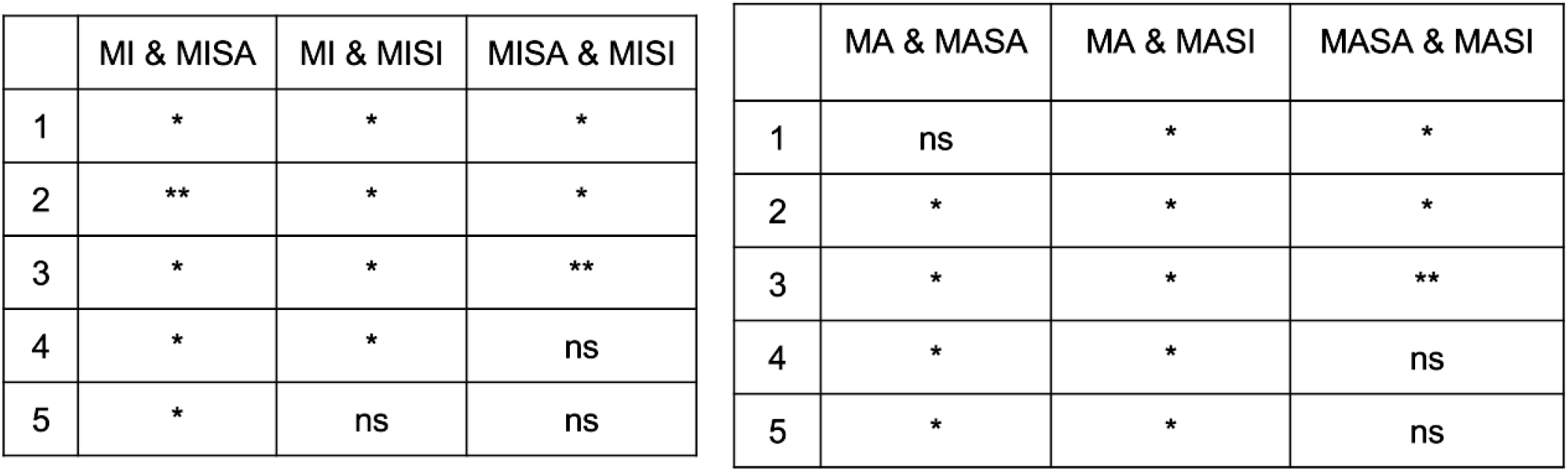
Significance measures for table S4

**Figure S6:**
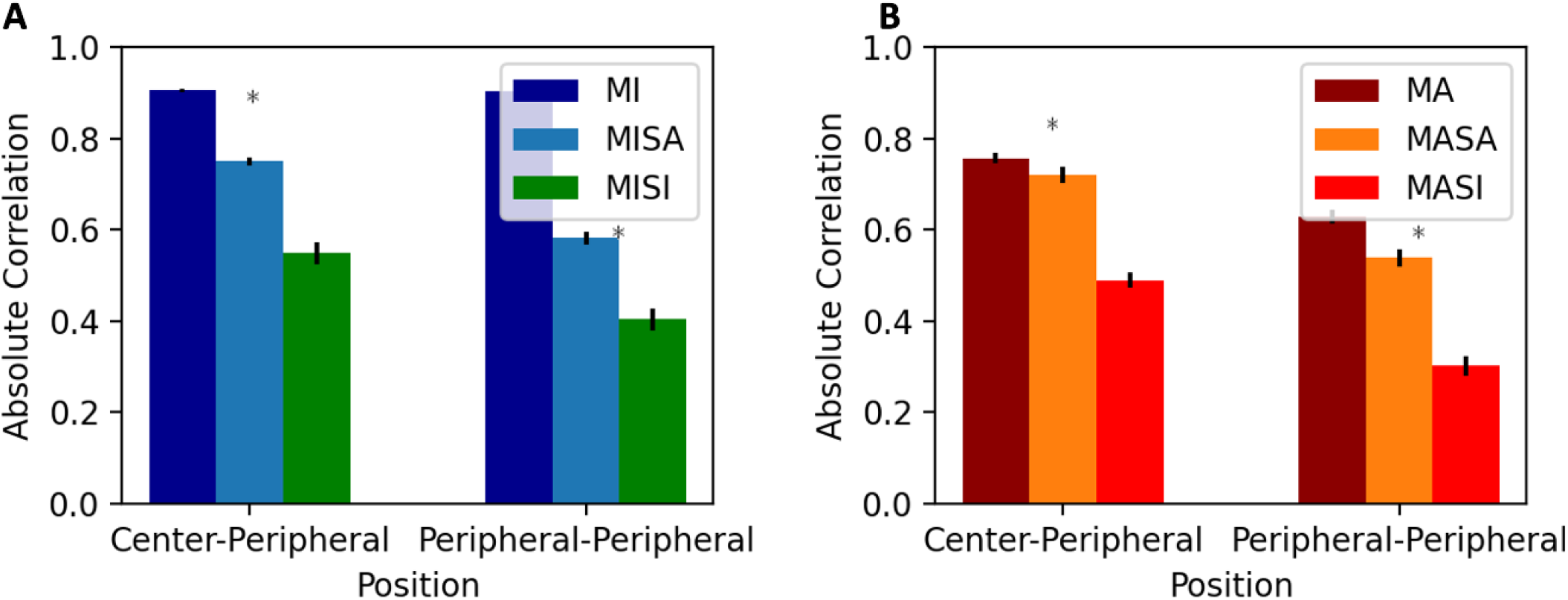
Parallel Connection. a) Variation of Absolute Correlation with addition of self-activation or self-inhibition to all the nodes of Mutual Inhibition based parallel connections. b) Variation of Absolute Correlation with the addition of self-activation or self-inhibition to all the nodes of Mutual Activation based parallel connections.

**Figure S7:**
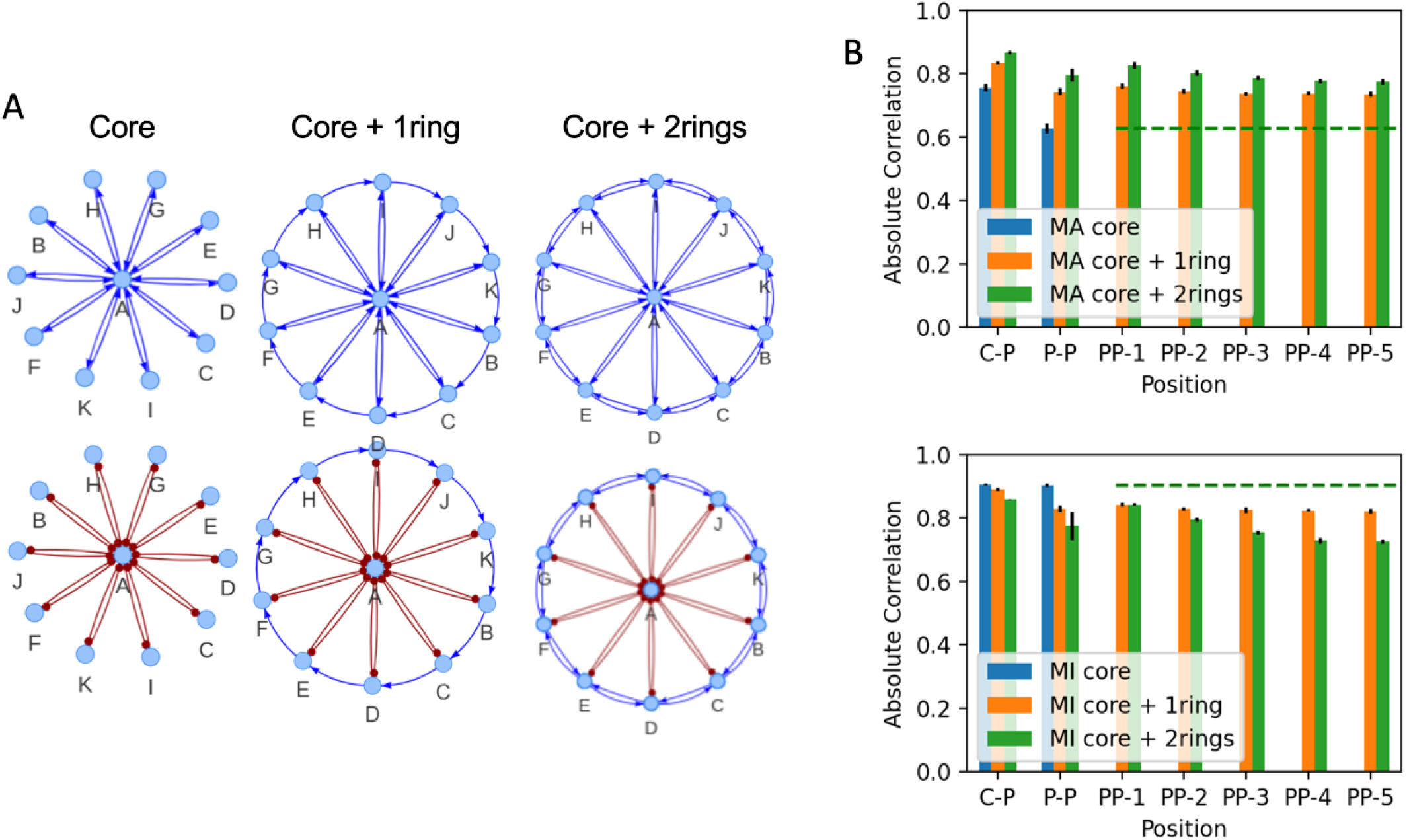
Addition of Mutual Activation rings to different cores. a) The network topology of the networks under consideration. MA and MI core are considered followed by addition of one activation and then mutual activation ring. b) Variation of Absolute Correlation with addition of activation ring and mutual activation ring to all nodes of MA and MI core connections.

**Figure S8:**
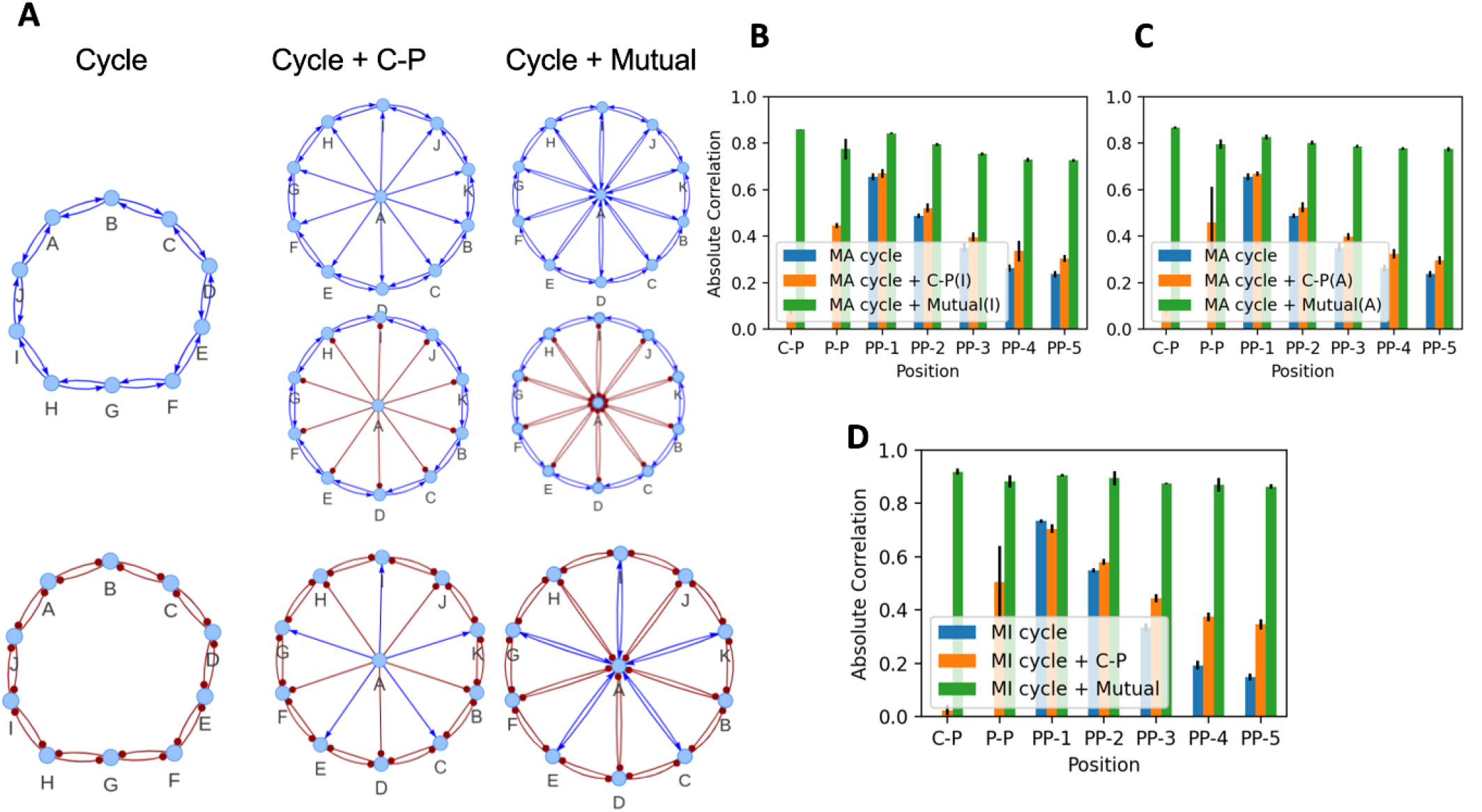
Addition of appropriate core to mutual activation and mutual inhibition rings. a) Mutual activation and mutual inhibition cores are added to mutual activation ring. Since the nature of stable states in of mutual inhibition ring is different, one possible configuration, that of a staggered core is added to the mutual inhibition ring. The corresponding network topologies are shown here. b) Variation of Absolute Correlation with addition of inhibition core to mutual activation ring. c) Variation of Absolute Correlation with addition of activation core to mutual activation ring. d) Variation of Absolute Correlation with addition of staggered core to mutual inhibition ring.

## References

[1] C. I. Abreu, V. L. Andersen Woltz, J. Friedman, and J. Gore. Microbial communities display alternative stable states in a fluctuating environment. PLOS Computational Biology, 16(5):1–17, 05 2020. doi: 10.1371/journal.pcbi.1007934. URL https://doi.org/10.1371/journal.pcbi.1007934.

[2] L. Agozzino, G. Balázsi, J. Wang, and K. A. Dill. How do cells adapt? stories told in landscapes. Annual Review of Chemical and Biomolecular Engineering, 11(1):155–182, 2020. doi: 10.1146/annurev-chembioeng-011720-103410. URL https://doi.org/10.1146/annurev-chembioeng-011720-103410. PMID: 32513086.

[3] D. Angeli, J. E. Ferrell, and E. D. Sontag. Detection of multistability, bifurcations, and hysteresis in a large class of biological positive-feedback systems. Proceedings of the National Academy of Sciences, 101(7):1822–1827, 2004. doi: 10.1073/pnas.0308265100. URL https://www.pnas.org/doi/abs/10.1073/pnas.0308265100.

[4] U. S. Bhalla, P. T. Ram, and R. Iyengar. Map kinase phosphatase as a locus of flexibility in a mitogen-activated protein kinase signaling network. Science, 297(5583):1018–1023, 2002. doi: 10.1126/science.1068873. URL https://www.science.org/doi/abs/10.1126/science.1068873.

[5] S. Bhattacharya, R. B. Conolly, N. E. Kaminski, R. S. Thomas, M. E. Andersen, and Q. Zhang. A Bistable Switch Underlying B-Cell Differentiation and Its Disruption by the Environmental Contaminant 2,3,7,8-Tetrachlorodibenzo-p-dioxin. Toxicological Sciences, 115(1):51–65, 02 2010. ISSN 1096-6080. doi: 10.1093/toxsci/kfq035. URL https://doi.org/10.1093/toxsci/kfq035.

[6] B.-S. Chen and C.-H. Wu. A systematic design method for robust synthetic biology to satisfy design specifications. BMC Systems Biology, 3(1):66, Jun 2009. ISSN 1752-0509. doi: 10.1186/1752-0509-3-66. URL https://doi.org/10.1186/1752-0509-3-66.

[7] B. Cummins, T. Gedeon, S. Harker, and K. Mischaikow. Dsgrn: Examining the dynamics of families of logical models. Frontiers in Physiology, 9, 2018. ISSN 1664-042X. doi: 10.3389/fphys.2018.00549. URL https://www.frontiersin.org/article/10.3389/fphys.2018.00549.

[8] S. Das and Debashis. Pulsatile signaling of bistable switches reveal the distinct nature of pulse processing by mutual activation and mutual inhibition loop. Journal of Theoretical Biology, 540:111075, 2022. ISSN 0022-5193. doi: https://doi.org/10.1016/j.jtbi.2022.111075. URL https://www.sciencedirect.com/science/article/pii/S002251932200073X.

[9] D. Del Vecchio, Y. Qian, R. M. Murray, and E. D. Sontag. Future systems and control research in synthetic biology. Annual Reviews in Control, 45:5–17, 2018. ISSN 1367-5788. doi: https://doi.org/10.1016/j.arcontrol.2018.04.007. URL https://www.sciencedirect.com/science/article/pii/S1367578818300191.

[10] A. Dey and D. Barik. Parallel arrangements of positive feedback loops limit cell-to-cell variability in differentiation. PLOS ONE, 12(11):1–20, 11 2017. doi: 10.1371/journal.pone.0188623. URL https://doi.org/10.1371/journal.pone.0188623.

[11] A. Dey and D. Barik. Potential landscapes, bifurcations, and robustness of tristable networks. ACS Synthetic Biology, 10(2):391–401, Feb. 2021. doi: 10.1021/acssynbio.0c00570. URL https://doi.org/10.1021/acssynbio.0c00570.

[12] A. Dhooge, W. Govaerts, and Y. A. Kuznetsov. Matcont: A matlab package for numerical bifurcation analysis of odes. ACM Trans. Math. Softw., 29(2):141–164, jun 2003. ISSN 0098-3500. doi: 10.1145/779359.779362. URL https://doi.org/10.1145/779359.779362.

[13] D. Dubnau and R. Losick. Bistability in bacteria. Molecular Microbiology, 61(3):564–572, 2006. doi: https://doi.org/10.1111/j.1365-2958.2006.05249.x. URL https://onlinelibrary.wiley.com/doi/abs/10.1111/j.1365-2958.2006.05249.x.

[14] A. S. Duddu, S. S. Majumdar, S. Sahoo, S. Jhunjhunwala, and M. K. Jolly. Emergent dynamics of a three-node regulatory network explain phenotypic switching and heterogeneity: a case study of th1/th2/th17 cell differentiation. Molecular Biology of the Cell, 0(0):mbc.E21–10–0521, 0. doi: 10.1091/mbc.E21-10-0521. URL https://doi.org/10.1091/mbc.E21-10-0521. PMID: 35353012.

[15] M. B. Elowitz and S. Leibler. A synthetic oscillatory network of transcriptional regulators. Nature, 403 (6767):335–338, Jan 2000. ISSN 1476-4687. doi: 10.1038/35002125. URL https://doi.org/10.1038/35002125.

[16] B. Errede, S. Hladyshau, N. Nivedita, D. Tsygankov, and T. C. Elston. Bistability in the polarity circuit of yeast. Mol Biol Cell, page mbcE20070445, May 2021.

[17] X. Fang, W. E. Bentley, and E. Zafiriou. Stochastic modeling of gene positive autoregulation networks involving signal molecules. Biophysical Journal, 95(7):3137–3145, 2008. ISSN 0006-3495. doi: https://doi.org/10.1529/biophysj.107.127175. URL https://www.sciencedirect.com/science/article/pii/S0006349508784583.

[18] P. C. Faucon, K. Pardee, R. M. Kumar, H. Li, Y.-H. Loh, and X. Wang. Gene networks of fully connected triads with complete auto-activation enable multistability and stepwise stochastic transitions. PLOS ONE, 9(7):1–16, 07 2014. doi: 10.1371/journal.pone.0102873. URL https://doi.org/10.1371/journal.pone.0102873.

[19] E. Feliu and C. Wiuf. Finding the positive feedback loops underlying multi-stationarity. BMC Systems Biology, 9(1):22, May 2015. ISSN 1752-0509. doi: 10.1186/s12918-015-0164-0. URL https://doi.org/10.1186/s12918-015-0164-0.

[20] J. E. Ferrell. Self-perpetuating states in signal transduction: positive feedback, double-negative feedback and bistability. Current Opinion in Cell Biology, 14(2):140–148, 2002. ISSN 0955-0674. doi: https://doi.org/10.1016/S0955-0674(02)00314-9. URL https://www.sciencedirect.com/science/article/pii/S0955067402003149.

[21] T. S. Gardner, C. R. Cantor, and J. J. Collins. Construction of a genetic toggle switch in escherichia coli. Nature, 403(6767):339–342, Jan 2000. ISSN 1476-4687. doi: 10.1038/35002131. URL https://doi.org/10.1038/35002131.

[22] S. Ghosh, K. Sureka, B. Ghosh, I. Bose, J. Basu, and M. Kundu. Phenotypic heterogeneity in mycobacterial stringent response. BMC Systems Biology, 5(1):18, Jan 2011. ISSN 1752-0509. doi: 10.1186/1752-0509-5-18. URL https://doi.org/10.1186/1752-0509-5-18.

[23] T. G. W. Graham, S. M. A. Tabei, A. R. Dinner, and I. Rebay. Modeling bistable cell-fate choices in the Drosophila eye: qualitative and quantitative perspectives. Development, 137(14):2265–2278, 07 2010. ISSN 0950-1991. doi: 10.1242/dev.044826. URL https://doi.org/10.1242/dev.044826.

[24] R. Guantes and J. F. Poyatos. Multistable decision switches for flexible control of epigenetic differentiation. PLOS Computational Biology, 4(11):1–13, 11 2008. doi: 10.1371/journal.pcbi.1000235. URL https://doi.org/10.1371/journal.pcbi.1000235.

[25] Z. Hamidouche, K. Rother, J. Przybilla, A. Krinner, D. Clay, L. Hopp, C. Fabian, A. Stolzing, H. Binder, P. Charbord, and J. Galle. Bistable Epigenetic States Explain Age-Dependent Decline in Mesenchymal Stem Cell Heterogeneity. Stem Cells, 35(3):694–704, 11 2016. ISSN 1066-5099. doi: 10.1002/stem.2514. URL https://doi.org/10.1002/stem.2514.

[26] K. Hari, B. Sabuwala, B. V. Subramani, C. A. M. La Porta, S. Zapperi, F. Font-Clos, and M. K. Jolly. Identifying inhibitors of epithelial–mesenchymal plasticity using a network topology-based approach. npj Systems Biology and Applications, 6(1):15, May 2020. ISSN 2056-7189. doi: 10.1038/s41540-020-0132-1. URL https://doi.org/10.1038/s41540-020-0132-1.

[27] K. Hari, P. Harlapur, A. Gopalan, V. Ullanat, A. S. Duddu, and M. K. Jolly. Emergent properties of coupled bistable switches. June 2021. doi: 10.1101/2021.06.15.448553. URL https://doi.org/10.1101/2021.06.15.448553.

[28] K. Hari, V. Ullanat, A. Balasubramanian, A. Gopalan, and M. K. Jolly. Landscape of epithelial mesenchymal plasticity as an emergent property of coordinated teams in regulatory networks. Dec. 2021. doi: 10.1101/2021.12.12.472090. URL https://doi.org/10.1101/2021.12.12.472090.

[29] S. Hati, A. S. Duddu, and M. K. Jolly. Operating principles of circular toggle polygons. Physical Biology, 18(4):046003, May 2021. doi: 10.1088/1478-3975/abef79. URL https://doi.org/10.1088/1478-3975/abef79.

[30] R. Hermsen, D. W. Erickson, and T. Hwa. Speed, sensitivity, and bistability in auto-activating signaling circuits. PLOS Computational Biology, 7(11):1–9, 11 2011. doi: 10.1371/journal.pcbi.1002265. URL https://doi.org/10.1371/journal.pcbi.1002265.

[31] B. Huang, M. Lu, D. Jia, E. Ben-Jacob, H. Levine, and J. N. Onuchic. Interrogating the topological robustness of gene regulatory circuits by randomization. PLOS Computational Biology, 13(3):1–21, 03 2017. doi: 10.1371/journal.pcbi.1005456. URL https://doi.org/10.1371/journal.pcbi.1005456.

[32] B. Huang, D. Jia, J. Feng, H. Levine, J. N. Onuchic, and M. Lu. Racipe: a computational tool for modeling gene regulatory circuits using randomization. BMC Systems Biology, 12(1):74, Jun 2018. ISSN 1752-0509. doi: 10.1186/s12918-018-0594-6. URL https://doi.org/10.1186/s12918-018-0594-6.

[33] N. T. Ingolia and A. W. Murray. Positive-feedback loops as a flexible biological module. Current Biology, 17(8):668–677, 2007. ISSN 0960-9822. doi: https://doi.org/10.1016/j.cub.2007.03.016. URL https://www.sciencedirect.com/science/article/pii/S0960982207011177.

[34] J. Jaruszewicz and T. Lipniacki. Toggle switch: noise determines the winning gene. Physical Biology, 10(3):035007, jun 2013. doi: 10.1088/1478-3975/10/3/035007. URL https://doi.org/10.1088/1478-3975/10/3/035007.

[35] D. Jia, M. K. Jolly, W. Harrison, M. Boareto, E. Ben-Jacob, and H. Levine. Operating principles of tristable circuits regulating cellular differentiation. Physical Biology, 14(3):035007, may 2017. doi: 10.1088/1478-3975/aa6f90. URL https://doi.org/10.1088/1478-3975/aa6f90.

[36] M. K. Jolly, M. Boareto, M. Lu, J. N. Onuchic, C. Clementi, and E. Ben-Jacob. Operating principles of notch–delta–jagged module of cell–cell communication. New Journal of Physics, 17(5):055021, may 2015. doi: 10.1088/1367-2630/17/5/055021. URL https://doi.org/10.1088/1367-2630/17/5/055021.

[37] M. Laurent and N. Kellershohn. Multistability: a major means of differentiation and evolution in biological systems. Trends in Biochemical Sciences, 24(11):418–422, 1999. ISSN 0968-0004. doi: https://doi.org/10.1016/S0968-0004(99)01473-5. URL https://www.sciencedirect.com/science/article/pii/S0968000499014735.

[38] J. Lee, J. Lee, K. S. Farquhar, J. Yun, C. A. Frankenberger, E. Bevilacqua, K. Yeung, E.-J. Kim, G. Balázsi, and M. R. Rosner. Network of mutually repressive metastasis regulators can promote cell heterogeneity and metastatic transitions. Proceedings of the National Academy of Sciences, 111(3): E364–E373, 2014. doi: 10.1073/pnas.1304840111. URL https://www.pnas.org/doi/abs/10.1073/pnas.1304840111.

[39] R. Lefever and W. Horsthemke. Bistability in fluctuating environments. implications in tumor immunology. Bulletin of Mathematical Biology, 41(4):469–490, 1979. ISSN 0092-8240. doi: https://doi.org/10.1016/S0092-8240(79)80003-8. URL https://www.sciencedirect.com/science/article/pii/S0092824079800038.

[40] C. Li and J. Wang. Landscape and flux reveal a new global view and physical quantification of mammalian cell cycle. Proceedings of the National Academy of Sciences, 111(39):14130–14135, Sept. 2014. doi: 10.1073/pnas.1408628111. URL https://doi.org/10.1073/pnas.1408628111.

[41] M. Lu, M. K. Jolly, R. Gomoto, B. Huang, J. Onuchic, and E. Ben-Jacob. Tristability in cancer-associated microrna-tf chimera toggle switch. The Journal of Physical Chemistry B, 117(42):13164–13174, Oct 2013. ISSN 1520-6106. doi: 10.1021/jp403156m. URL https://doi.org/10.1021/jp403156m.

[42] P. Maheshwari and R. Albert. A framework to find the logic backbone of a biological network. BMC Systems Biology, 11(1):122, Dec 2017. ISSN 1752-0509. doi: 10.1186/s12918-017-0482-5. URL https://doi.org/10.1186/s12918-017-0482-5.

[43] S. Matsuoka and M. Ueda. Mutual inhibition between pten and pip3 generates bistability for polarity in motile cells. Nature Communications, 9(1):4481, Oct 2018. ISSN 2041-1723. doi: 10.1038/s41467-018-06856-0. URL https://doi.org/10.1038/s41467-018-06856-0.

[44] E. M. Ozbudak, M. Thattai, H. N. Lim, B. I. Shraiman, and A. van Oudenaarden. Multistability in the lactose utilization network of escherichia coli. Nature, 427(6976):737–740, Feb 2004. ISSN 1476-4687. doi: 10.1038/nature02298. URL https://doi.org/10.1038/nature02298.

[45] R. Perez-Carrasco, C. P. Barnes, Y. Schaerli, M. Isalan, J. Briscoe, and K. M. Page. Combining a toggle switch and a repressilator within the ac-dc circuit generates distinct dynamical behaviors. Cell Systems, 6(4):521–530.e3, 2018. ISSN 2405-4712. doi: https://doi.org/10.1016/j.cels.2018.02.008. URL https://www.sciencedirect.com/science/article/pii/S2405471218300619.

[46] R. Planqué, F. J. Bruggeman, B. Teusink, and J. Hulshof. Understanding bistability in yeast glycolysis using general properties of metabolic pathways. Mathematical Biosciences, 255:33–42, 2014. ISSN 0025-5564. doi: https://doi.org/10.1016/j.mbs.2014.06.006. URL https://www.sciencedirect.com/science/article/pii/S0025556414001126.

[47] A. Randall, P. Guye, S. Gupta, X. Duportet, and R. Weiss. Chapter seven - design and connection of robust genetic circuits. In C. Voigt, editor, Synthetic Biology, Part A, volume 497 of Methods in Enzymology, pages 159–186. Academic Press, 2011. doi: https://doi.org/10.1016/B978-0-12-385075-1.00007-X. URL https://www.sciencedirect.com/science/article/pii/B978012385075100007X.

[48] E. R. Regan and W. C. Aird. Dynamical systems approach to endothelial heterogeneity. Circulation Research, 111(1):110–130, 2012. doi: 10.1161/CIRCRESAHA.111.261701.

[49] D. Sprinzak, A. Lakhanpal, L. LeBon, L. A. Santat, M. E. Fontes, G. A. Anderson, J. Garcia-Ojalvo, and M. B. Elowitz. Cis-interactions between notch and delta generate mutually exclusive signalling states. Nature, 465(7294):86–90, Apr. 2010. doi: 10.1038/nature08959. URL https://doi.org/10.1038/nature08959.

[50] D. Stockholm, F. Edom-Vovard, S. Coutant, P. Sanatine, Y. Yamagata, G. Corre, L. Le Guillou, T. M. A. Neildez-Nguyen, and A. Pàldi. Bistable cell fate specification as a result of stochastic fluctuations and collective spatial cell behaviour. PLOS ONE, 5(12):1–12, 12 2011. doi: 10.1371/journal.pone.0014441. URL https://doi.org/10.1371/journal.pone.0014441.

[51] D. Stockholm, F. Edom-Vovard, S. Coutant, P. Sanatine, Y. Yamagata, G. Corre, L. Le Guillou, T. M. A. Neildez-Nguyen, and A. Pàldi. Bistable cell fate specification as a result of stochastic fluctuations and collective spatial cell behaviour. PLOS ONE, 5(12):1–12, 12 2011. doi: 10.1371/journal.pone.0014441. URL https://doi.org/10.1371/journal.pone.0014441.

[52] T. Tian and K. Burrage. Stochastic models for regulatory networks of the genetic toggle switch. Proceedings of the National Academy of Sciences, 103(22):8372–8377, 2006. doi: 10.1073/pnas.0507818103. URL https://www.pnas.org/doi/abs/10.1073/pnas.0507818103.

[53] T.-L. To and N. Maheshri. Noise can induce bimodality in positive transcriptional feedback loops without bistability. Science, 327(5969):1142–1145, 2010. doi: 10.1126/science.1178962. URL https://www.science.org/doi/abs/10.1126/science.1178962.

[54] J.-W. Veening, W. K. Smits, and O. P. Kuipers. Bistability, epigenetics, and bet-hedging in bacteria. Annual Review of Microbiology, 62(1):193–210, 2008. doi: 10.1146/annurev.micro.62.081307.163002. URL https://doi.org/10.1146/annurev.micro.62.081307.163002. PMID: 18537474.

[55] J. Wang, L. Xu, and E. Wang. Potential landscape and flux framework of nonequilibrium networks: Robustness, dissipation, and coherence of biochemical oscillations. Proceedings of the National Academy of Sciences, 105(34):12271–12276, Aug. 2008. doi: 10.1073/pnas.0800579105. URL https://doi.org/10.1073/pnas.0800579105.

[56] L. Wang, B. L. Walker, S. Iannaccone, D. Bhatt, P. J. Kennedy, and W. T. Tse. Bistable switches control memory and plasticity in cellular differentiation. Proceedings of the National Academy of Sciences, 106(16):6638–6643, 2009. doi: 10.1073/pnas.0806137106. URL https://www.pnas.org/doi/abs/10.1073/pnas.0806137106.

[57] F. Wu, R.-Q. Su, Y.-C. Lai, and X. Wang. Engineering of a synthetic quadrastable gene network to approach waddington landscape and cell fate determination. eLife, 6:e23702, apr 2017. ISSN 2050-084X. doi: 10.7554/eLife.23702. URL https://doi.org/10.7554/eLife.23702.

[58] Y. Yang, J. L. Nemhauser, and E. Klavins. Synthetic bistability and differentiation in yeast. ACS Synthetic Biology, 8(5):929–936, May 2019. doi: 10.1021/acssynbio.8b00524. URL https://doi.org/10.1021/acssynbio.8b00524.

[59] T.-M. Yi, Y. Huang, M. I. Simon, and J. Doyle. Robust perfect adaptation in bacterial chemotaxis through integral feedback control. Proceedings of the National Academy of Sciences, 97(9):4649–4653, 2000. doi: 10.1073/pnas.97.9.4649. URL https://www.pnas.org/doi/abs/10.1073/pnas.97.9.4649.

[60] R. Zhu, J. M. del Rio-Salgado, J. Garcia-Ojalvo, and M. B. Elowitz. Synthetic multistability in mammalian cells. Science, 375(6578):eabg9765, 2022. doi: 10.1126/science.abg9765. URL https://www.science.org/doi/abs/10.1126/science.abg9765.

